# The immunodominant protein P116 extracts cholesterol and other essential lipids

**DOI:** 10.1101/2022.06.28.498018

**Authors:** Lasse Sprankel, David Vizarraga, Jesús Martín, Sina Manger, Jakob Meier-Credo, Marina Marcos, Josep Julve, Noemi Rotllan, Margot P. Scheffer, Joan Carles Escolà-Gil, Julian D. Langer, Jaume Piñol, Ignacio Fita, Achilleas S. Frangakis

## Abstract

Many human pathogens need to extract lipids from their environment for survival and proliferation. How this is accomplished on a molecular level is largely unknown^1^. Here, we report a comprehensive structural and functional analysis of the previously uncharacterized protein P116 (MPN_213) from *Mycoplasma pneumoniae*, a human pathogen responsible for approximately 30% of community-acquired human pneumonia^2^. Single-particle cryo-electron microscopy of P116 at 3.3 Å resolution reveals a homodimer with a core domain presenting a previously unseen fold. This fold creates a large cavity of ∼18,000 Å^3^ with a hydrophobic internal surface that is accessible to solvent. Within the cavity ligands with a length of 10-19 Å and a width of 4 Å could be observed. These ligands were identified as the essential lipids phosphatidylcholine, sphingomyelin and cholesterol using mass spectrometry. When the cavity is emptied, the protein undergoes an extensive conformational change that can no longer accommodate lipids. When emptied P116 is incubated with high-density lipoproteins (HDLs) a net transfer of cholesterol is demonstrated by a radioactivity experiment and cryo-electron microscopy resolves a complex between P116 and HDL. Taken together, our results reveal the mechanism by which P116 extracts essential lipids from the host environment and possibly then delivers them into the membrane by a wringing movement. This mechanism may be precedential for other cholesterol-auxotrophic bacteria.

## Introduction

*Mycoplasma pneumoniae* is a facultative intracellular human pathogen causing community-acquired pneumonia that can manifest severe systemic effects^2^. Unlike other respiratory pathogens, *M. pneumoniae* has no approved vaccine^3^. *Mycoplasmas* lack a cell wall and have the smallest known genomes^4^. *M. pneumoniae*, with a 816 kb genome, is a model organism for a minimal cell^5^. Many of the metabolic pathways required to synthesize essential products are absent, which makes an uptake by specialized mechanisms necessary. In fact, *M. pneumoniae* cannot synthesize several of the lipids that are important components of the cell membrane, such as sphingomyelin, phosphatidylcholine and cholesterol^6^. Instead, it must take up lipids from the host environment and adapts its membrane composition depending on the medium in vitro^*7–9*^. Cholesterol in particular, which is present in only a few prokaryotes including the clinically relevant *Helicobacter pylori* and *Borrelia burgdorferi*, is essential for *M. pneumoniae* cells and is the most abundant lipid in the membranes, accounting for 35–50% of the total lipid fraction^7^. To date, it is unclear how Mycoplasma and other prokaryotic species achieve lipid uptake from the environment.

In this work, we report the structural and functional characterization of P116, a strongly immunogenic and essential protein for the viability of *M. pneumoniae* cells. P116 was previously uncharacterized, although it has been reported to potentially contribute to adhesion to host cells^10^. Despite the essential role of P116 the *M. pneumoniae* genome contains only a single copy of *p116* (*mpn213*). To elucidate the role of P116, we first determined the structure of the ectodomain by single-particle cryo-electron microscopy (cryoEM). The structure has a novel fold (with no matches in the Protein Data Bank) featuring a uniquely large hydrophobic cavity that is fully accessible to solvent. Mass spectrometry and other analytical techniques identify ligands found in the cavity as several different lipids (incl. cholesterol), some of which are essential. Based on these findings, we describe the mechanism by which *Mycoplasmas* can extract lipids from the environment and possibly also deposit them in their own membrane, thus explaining the essential role of P116 for the survival of *M. pneumoniae* cells.

## Results

### P116 is abundant on the cell surface

A construct predicted to span the whole ectodomain of P116 from *M. pneumoniae* (residues 30–957) was overexpressed in *Escherichia coli* and purified by His-tag affinity and gel filtration chromatography (Materials and Methods and **Extended Data Figure 1**). Immunolabeling with both polyclonal and monoclonal antibodies against this construct showed an intense and uniform distribution of labeling across the whole surface of *M. pneumoniae* cells (**Figure 1a**), with adhesion and motility unaffected by the antibodies. This distribution contrasts with that of P1, an adhesion protein that concentrates at the tip of the cell and has strong effects on adhesion and motility^11,12^.

**Figure 1.**
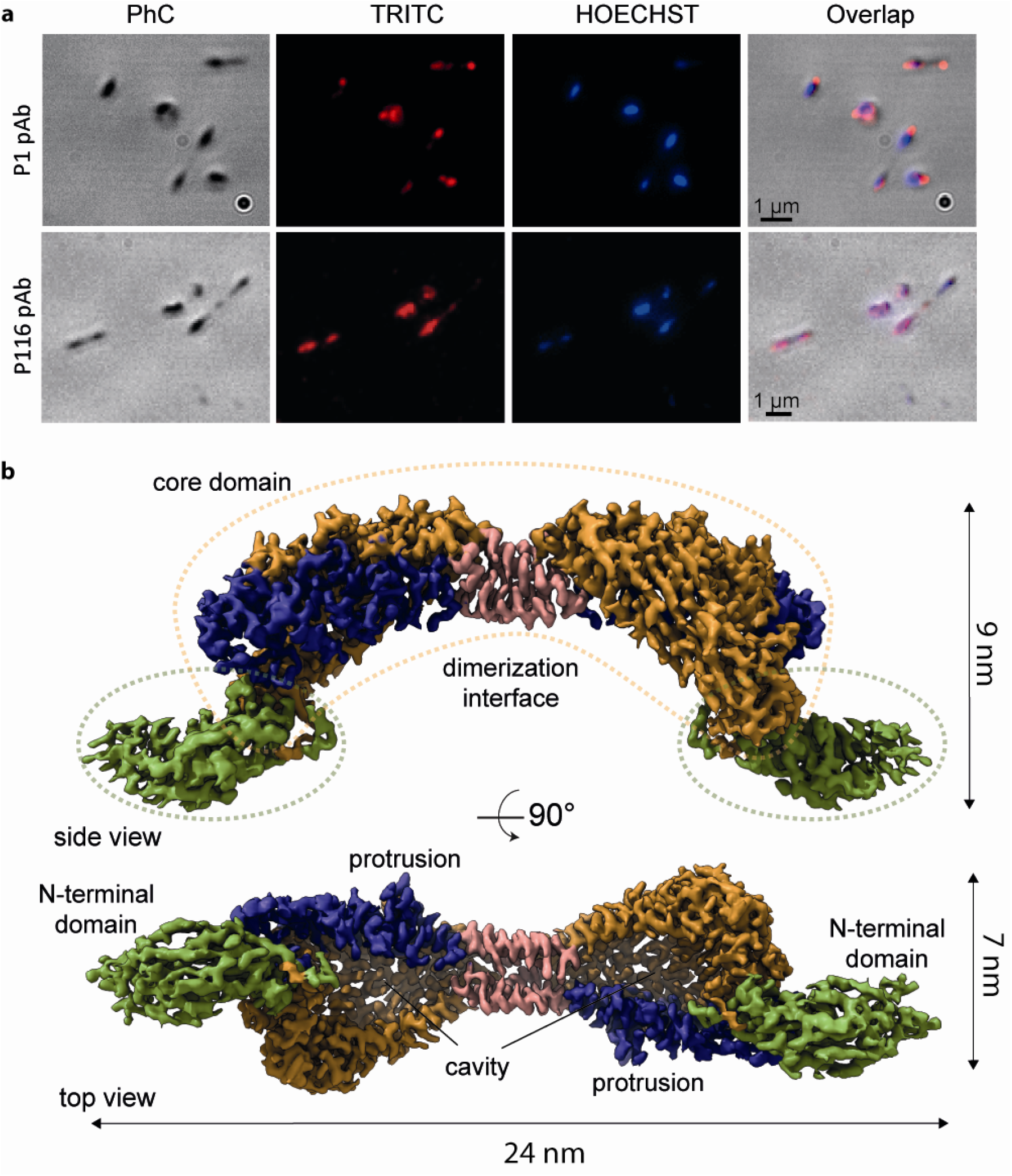
Structure of P116 and its localization in *Mycoplasma pneumoniae* cells. **a)** Phase contrast (PhC) immunofluorescence microscopy images *of M. pneumoniae* cells using labeling with polyclonal antibodies against the ectodomains of adhesin P1 (top row; used as a reference) and P116 (bottom row). Labelling for P1 concentrates at the tip of the cell, while for P116 it covers the whole surface homogenously. **b)** Two views of the cryoEM density map of the complete extracellular region of the P116 dimer at 3.3 A□ resolution, 90 degrees apart. The homodimer is held together by the dimerization interface (shown in pink). The core domains have four contiguous antiparallel helices (shown in blue) and a β-sheet with five antiparallel strands (shown in orange). The N-terminal domain is shown in green. The top view displays a huge cavity that is fully accessible to solvent. The cleft providing access to the cavity spans the whole core domain. Each monomer also has a distinct protrusion (shown in blue as part of the antiparallel α-helices).

### P116 has a novel fold with a lipid-accessible cavity

The structure of P116 (30–957) was determined by single-particle cryoEM at 3.3 A□ resolution (according to the gold standard criterion of FSC 0.143; **Extended Data Table I, Extended Data Figure 2**. It is an elongated homodimer of ∼240 Å along its longest axis, which adopts an arched shape with an arc-radius of 200 Å (**Figure 1b, Extended Data Movies 1, 2**). Each monomer consists of two distinct subunits: A N-terminal domain (residues 60–245), situated distal to the dimer axis, and a core domain (residues 246–867). Proximal to the dimer axis is the dimerization interface (**Figure 1b, Extended Data Figure 3**), which is very well resolved. In addition, the N-terminal domain has significant hinge mobility with respect to the core domain, which reduced the local resolution of the cryoEM map (**Extended Data Figure 2**), making model building difficult for the most distal parts of the construct (see Materials and Methods and **Extended Data Figure 4**). The homodimer displays significant flexibility with many vibrational modes, as classification illustrates (**Extended Data Figure 5**). Finally, some residues at the N- and C-termini of the construct (30–59 and 868–957, respectively) were not visible in the cryoEM maps. The flexibility of the homodimer involves a change in the curvature of approximately 100 Å, wringing along the axis perpendicular to the dimer axis by ∼80 degrees, and bending up to 20 degrees (**Extended Data Figure 5, Extended Data Movie 3**).

The core domain resembles a half-opened left hand, with four contiguous antiparallel α-helices corresponding to the four fingers and the N-terminal domain the thumb (**Figure 2a**). The helices corresponding to the wrist form the dimer interface, and a conserved tryptophan residue (Trp681) interacts tightly with the neighboring monomer. In the variant Trp681Ala, the rate of dimers to monomers is 1:4, compared to only dimers without the mutation (**Extended Data Figure 3b**). The palm of the hand includes a long and well defined central α-helix, the bridge helix (residues 268– 304), and a rigid β-sheet of five antiparallel strands that extends to the N-terminal domain (**Figure 2b**). The hand appears in a half-opened state with a large elongated cleft across the whole core domain (**Figure 2c**). The inner part of the hand (i.e. the fingers and palm) forms a large cavity that measures 62 Å proximal to distal and 38 Å anterior to posterior with a volume of ∼18,000 Å^3^. The cavity is completely hydrophobic although fully accessible to the solvent (**Figure 2c, Extended Data Movie 4)**. In addition, the core has two access points, one at the dorsal side and one at the distal side (**Figure 3a**). Using the DALI server, we found only very weak structural relationships between P116 and all other experimentally determined protein structures available in the Protein Data Bank, which shows that P116 has a new, unique fold.

**Figure 2.**
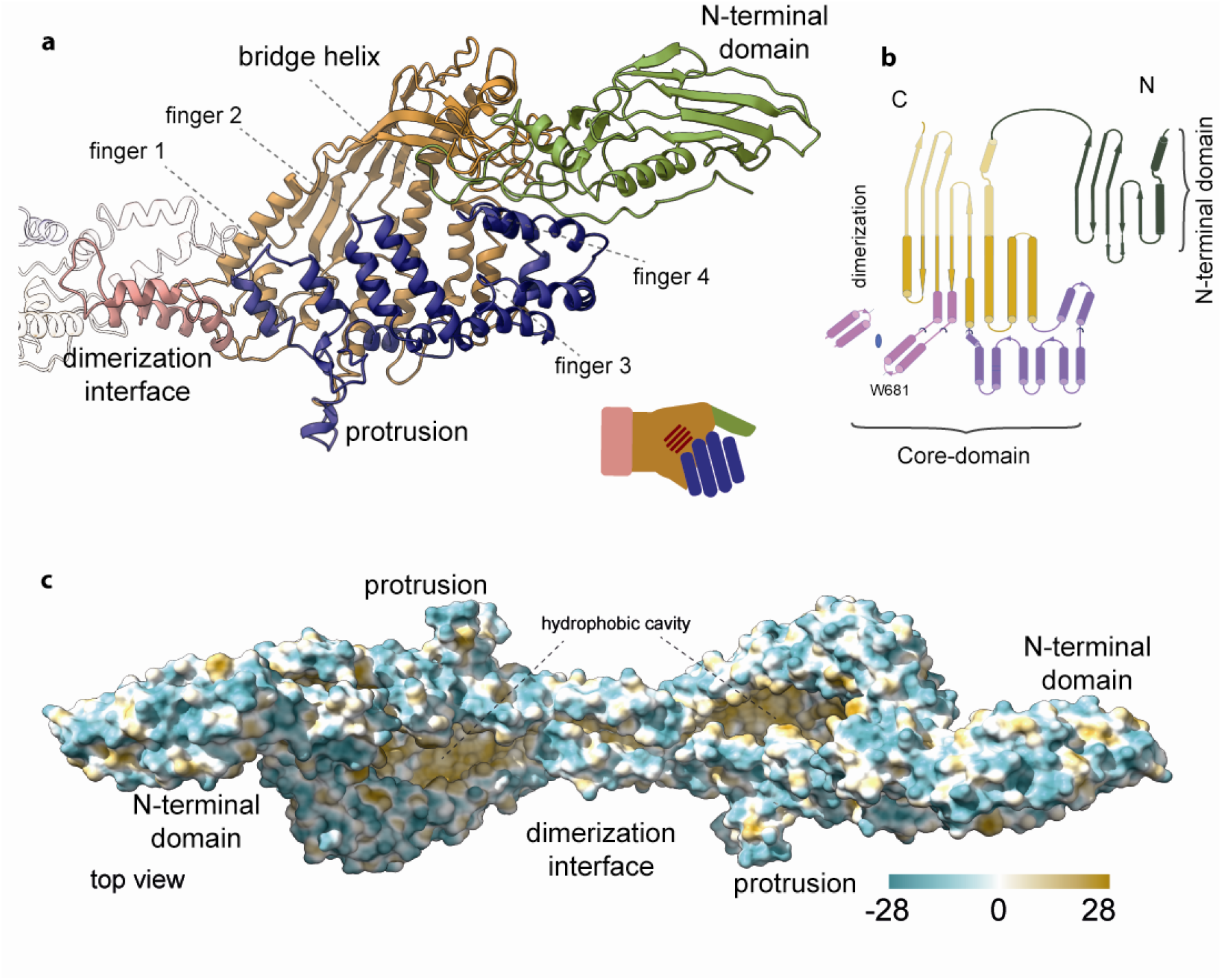
P116 structure and hydrophobic areas. **a)** Ribbon model of the P116 monomer, colored as in Fig. 1. The overall shape of the structure corresponds to a left hand, with the four antiparallel α-helices representing fingers (shown in blue), and the bridge helix and β-sheet of five antiparallel strands representing the palm. The N-terminal domain, which is very flexible, corresponds to the thumb. The dimerization helices (shown in pink) correspond to the wrist. **b)** The overall topology of P116. The N-terminal and core domains of P116 share a similar topology, which suggests that P116 might have been generated by duplication of an ancestor domain. **c)** The hydrophobic map of the P116 homodimer shows that the cavity in the core domain is hydrophobic.

**Figure 3:**
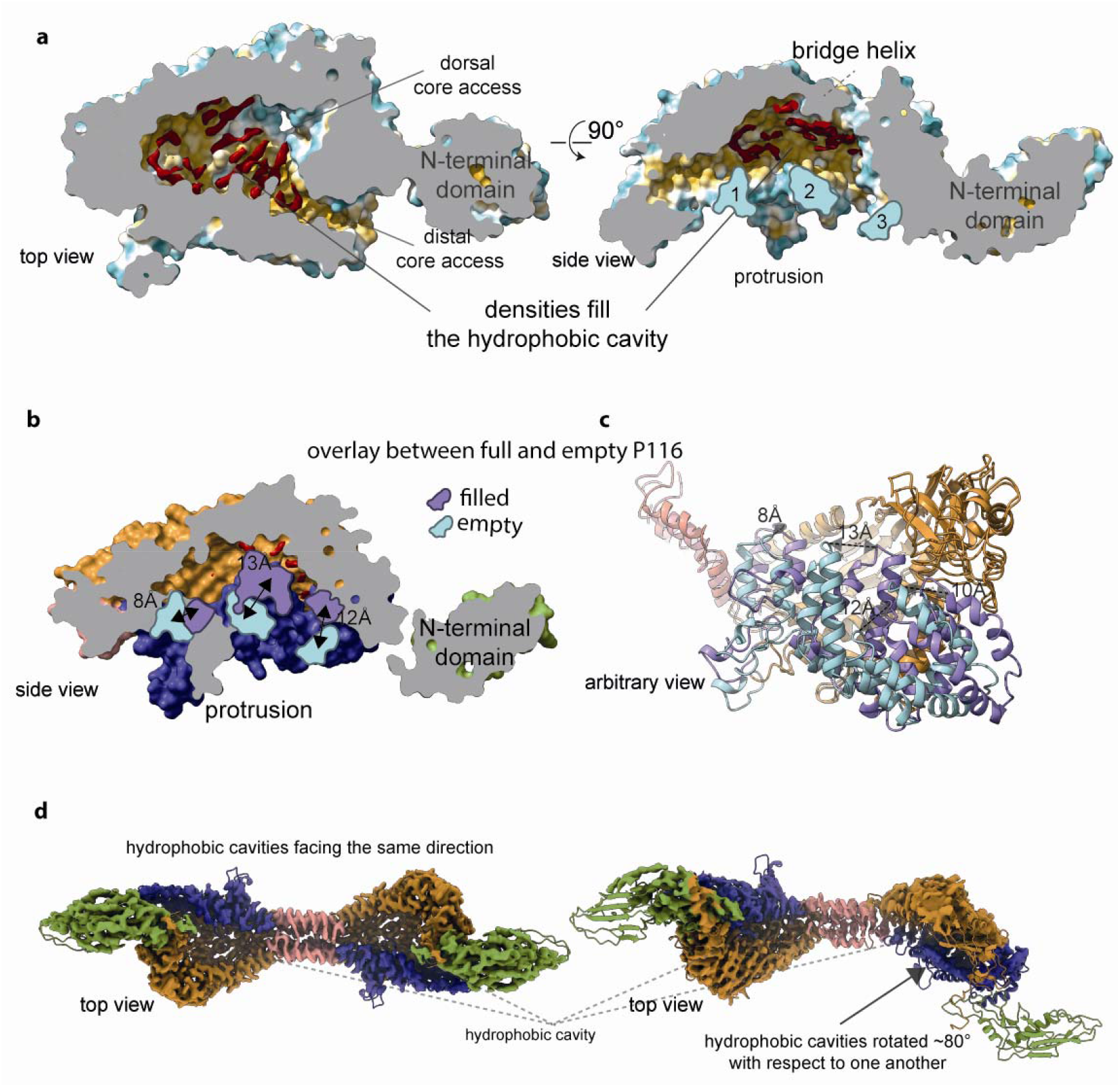
Purified P116 is filled with ligands and displays a large conformational variation compared to empty P116. **a)** Cross-section through the core domain of original P116 exposes a series of elongated densities (shown in red), which cannot be accounted for by the structure. These densities are ∼4 Å wide and 10–19 Å long and are surrounded by highly conserved hydrophobic residues. The cross-section also reveals that the core domain can be accessed dorsally and distally. The side view of the core domain shows that the densities are aligned to the bridge helix and away from the fingers (shown in red). The individual fingers are indicated with digits 1 to 3 (finger 4 is not visible in this illustration). **b)** Overlay between empty and full P116. Side view of the cross-section surface view of the empty and full P116 shows that the fingers (in purple) have come closer to the core domain, massively reducing the available volume. Their new position is markedly different compared to the full P116 (shown in light blue). Finger 1 moved 8Å sideways and towards the core, finger 2 has moved 13Å towards the core and Finger 3 has moved 12Å towards the core. The volume in the empty P116 is not sufficient to accommodate ligands anymore. **c)** In the ribbon presentation the conformation differences between the empty and full P116 structures can be seen. All four fingers (antiparallel α-helices) have moved towards the inner part of the hand (individual distances are indicated filled conformation in light blue, empty conformation in purple). **d)** Two cryoEM classes reveal a wringing movement of P116. Comparison of the two density maps (superimposed with the ribbon diagram of the structure) shows that the wringing movement of P116 allows for the two hydrophobic cavities in the dimer to face almost opposite directions. The top view on the left shows both cavities facing in one direction, while the top view on the right shows the cavities rotated ∼80 degrees to each other.

The N-terminal domain is compact and organized around a cluster of aromatic residues, at the center of which is the only tryptophan residue of the domain (Trp121). The N-terminal and core domains of P116 superimpose for 126 equivalent residues (68% of the N-terminal domain), suggesting that P116 might have been generated by duplication of an ancestor domain. The common secondary structural elements in the N-terminal and core domains consist of a β-sheet and the two helices preceding the sheet (**Figure 2b**). The core domain is much larger than the N-terminal domain mainly due to two insertions containing twelve and four helices, respectively.

For the inner part of the P116 core domain, the cryoEM maps show prominent elongated densities (with a length of 10–19 Å and a width of 4 Å) that fill most of the hydrophobic areas (**Figure 3a, Extended Data Movies 5, 6**). These elongated densities, which are unaccounted for, cannot be explained by the protein residues missing in the model. Instead, the mass excess of ∼13 kDa, consistently measured by multiple angle light scattering (MALS) and mass spectrometry for P116 in different preparations, could be explained by the presence of ligand molecules bound to P116 (**Figure 4a**). Initial mass spectrometry analysis of the same samples from which the structure of P116 was determined (see Materials and Methods) showed the presence of several lipid species, including phosphatidylcholine and sphingomyelin, which are essential for *M. pneumoniae*^3^, and of wax esters (**Figure 4b and Extended Data Figure 6**).

**Figure 4:**
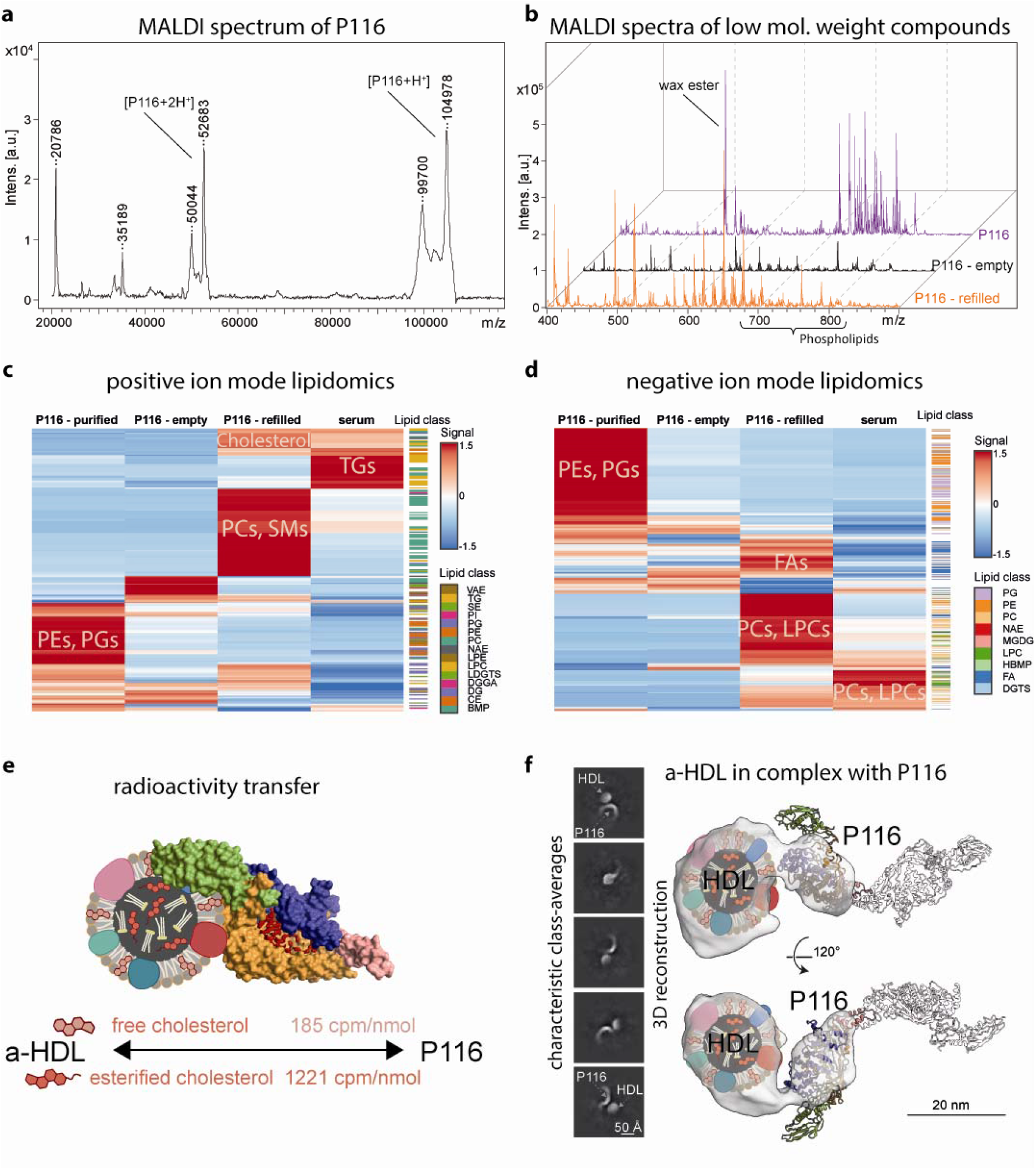
**a)** MALDI-TOF mass spectrum of original P116 sample (linear mode, high mass range), showing a dominant peak at the 105 kDa corresponding to the singly charged full protein, as well as the charges states two, three and four. **b)** Stacked MALDI-TOF mass spectra (reflector mode, low mass range) of the originally purified P116 (purple, back), the empty P116 (black, middle) and the refilled P116 sample (orange, front) showing a change in the lipid distribution among the samples. **c**) and **d**) Hierarchical clustering of lipid compounds identified in positive (c) and negative (d) ion mode lipidomics (LC-ESI-IMS-MS/MS) analyses, showing differential distributions of lipid compositions in original P116 (first column), emptied P116 (second column), refilled P116 (third column) and serum (fourth column), respectively. All data were normalized to the mTIC of all identified compounds in each sample and row-wise scaling was applied. **e)** When radiolabeled HDLs (here presented schematically) are incubated with P116, a net cholesterol transfer to P116 can be measured as indicated by the number at the flux arrow (for both free and esterified cholesterol). **f)** CryoEM analysis of empty P116 incubated with HDL shows that P116 binds HDLs between its N-terminal and core domains and is refilled. P116 is attached to HDL through its distal core access. Due to the flexibility of P116 and the variability of HDL, only one subunit of P116 can be seen at this threshold. Reducing the threshold causes the second subunit to appear.

Among the P116 orthologues found in a BlastP search^13^ of eight *Mycoplasma* species, the amino acids lining the hydrophobic cavity are largely conserved (either identical or with similar characteristics), in particular those in the proximity of the unaccounted, elongated densities (**Extended Data Figure 7**). Modeling P116 and its orthologues with AlphaFold^14^ results in all the models having a similar tertiary structure, in which a large core domain is flanked by a smaller N-terminal domain, but the relative position of the domains does not closely match the experimental structure.

### The conformation of empty P116 cannot accommodate lipid binding

To obtain ‘empty’ P116 that was free of any bound ligands, we treated the P116 samples with the detergent Triton-X 100 (see below and Materials and Methods). Mass spectrometry confirmed a massive reduction of lipids in the sample (**Figure 4b)**. The structure of the empty P116 sample was solved by cryoEM at 4 Å resolution (**Extended Data Figure 8**). Its overall topology is almost identical to that of the original P116 sample, with the difference that the cavity is closed as a result of fingers 1, 2 and 3 being closer to the palm by 8, 13 and 12 Å, respectively, and finger 4 moving 11 Å sideways to retain the distal core access to the palm (**Figure 3b, Extended Data Movies 7, 8**, and **Extended Data Figure 9**). These changes reduce the volume within the core domain from ∼18,000 Å^3^ to ∼6,300 Å^3^. The unoccupied volume between the fingers and palm reduces to two pockets that are large enough for lipids to pass through but appear unoccupied in the cryoEM density. A comparison of the filled and empty P116 structures shows that the original densities that were unaccounted for create massive steric clashes in the closed configuration of the fingers, demonstrating that the cavity can no longer accommodate lipids (**Extended Data Movie 9**). In the empty P116, the dimerization interface is shifted towards the dorsal side of the molecule by 10 Å, resulting in a contraction that changes the arc radius of the dimer from 500 to 600 Å and shifts the N-terminal domain towards the dimerization interface.

### Refilled P116 is structurally identical to the purified sample

We next refilled the empty P116 samples by incubating them either with fetal bovine serum (FBS) or with high-density lipoproteins (HDL) and then re-purified them by affinity chromatography. Media containing FBS is a common growing broth for *M. pneumoniae* cultures, although lipoproteins, in particular HDL, are efficient substitutes for serum in mycoplasma culture media, likely because lipoproteins can provide the key lipids, in particular cholesterol, which is essential for mycoplasma cells^15^. We solved the structure of the refilled P116 samples at 3.5 Å resolution using cryoEM. The structure of the refilled P116 is practically identical at 3.5 Å resolution to the structure of the original P116 sample, including densities at the palm of the hand that can be assigned to ligands. Mass spectrometry of the refilled samples shows the clear presence of lipids (**Figure 4b**). Classes of subunits of the dimer show a wringing of ∼80 degrees (**Figure 3d, Extended Data Figure 5** and **Extended Data Movie 3**).

### P116 is conformationally flexible

In the original P116, empty P116 and refilled P116 samples, the structure appears predominantly as a homodimer. In all cases, the homodimer exhibits significant flexibility. Most prominently, the empty structure has a different arc radius than those of the original and refilled structures. In the original and refilled structures, a wringing motion is visible: each monomer is twisted in the opposite direction along the axis perpendicular to the dimer axis (**Figure 3d, Extended Data Movie 3** and **Extended Data Figure 5**). In all P116 structures, the N-terminal domain is the most flexible. Within the core domain, temperature factors are higher at the fingertips, indicating the movement of the antiparallel α-helices. When the fingers approach the palm, this results in a closing of the hand and a clash with the densities therein (**Extended Data Movie 9**).

### P116 ligands include essential lipids

We next set out to characterize the possible ligands within P116. We first measured the rate of radioactivity transfer to P116 after incubation with HDL particles containing either tritium-labeled cholesterol ([^3^H]cholesterol) or tritium-labeled cholesteryl oleate as a representative of cholesterol esters (**Table I**). A significant fraction of the HDL-[^3^H]-radiotracer was detected in the post-incubated and purified P116 fractions, indicating a net transfer of both cholesterol and cholesterol ester between HDL and P116. The total absence of the most abundant HDL protein (APOA1), cross-checked by immune detection, verified that no HDL remnants had contaminated the purified P116 fractions. The highest rate of radiotracer transfer was achieved when [^3^H]cholesterol-containing HDLs were mixed with empty P116. Transfer of [^3^H]cholesterol was also present, although reduced, when the original P116 was incubated with labeled HDL. Transfer of [^3^H]cholesterol esters to P116 would require a direct interaction between HDL and P116, as these esters are buried in the core of the HDL particles (**Table I**). Passive cholesterol transport has been reported from cellular membranes to HDL or from LDL to HDL^16^, but the concept that bacteria can exploit such a mechanism is completely new. The net flux of cholesterol is bidirectional and is governed by the cholesterol gradient between acceptor and donor molecules.

**Table I:** Relative transfer of (esterified) cholesterol from HDL to P116.

|  | % of<br>[<sup>3</sup>H]cholesterol<br>transferred/mL | nmol<br>cholesterol<br>transferred/mL/h | nmol cholesterol<br>transferred/mg<br>P116<br>* |
| --- | --- | --- | --- |
| **HDL to empty P116** |  |  |  |
| **Free cholesterol** | 13.12 | 13.52 | 59.49 (6.3) |
| **Esterified cholesterol** | 6.98 | 7.22 | 31.75 (3.3) |
| **HDL to original P116** |  |  |  |
| **Free cholesterol** | 7.89 | 7.42 | 32.63 (3.4) |
| **Esterified cholesterol** | 6.32 | 6.01 | 26.44 (2.8) |

\* Numbers in parentheses are the estimated number of cholesterol molecules transferred per P116 subunit (assuming a Mw of ~105 KDa for the construct).

We then conducted a detailed liquid chromatography-coupled mass spectrometry (LC-MS) analysis. We identified more than 500 lipid species in the samples and found striking differences between the original, empty and refilled P116 samples (**Figure 4c, 4d**). Characterization of the lipids in the original and refilled P116 samples showed the presence of phosphatidylcholine and sphingomyelin lipids, among others, which are essential for *M. pneumoniae*. While these analyses found wax esters in the original P116, far fewer were found in the refilled P116. Wax esters are not known to be required by *M. pneumoniae*, although some pathogenic bacteria use wax esters as a carbon source^17,18^. However, wax esters are part of the cultivation medium of the *E. coli* strain in which P116 was produced. These findings are in agreement with the fact that *M. pneumoniae* takes up and incorporates many lipid species and adapts its membrane composition to the available lipid spectrum. In the P116 samples refilled from FBS, a clear accumulation of the essential lipids phosphatidylcholine and sphingomyelin, as well as cholesterol molecules, can be seen (**Figure 4c, 4d** and **Extended Data Table II**). These findings are in strong agreement with the functional data from the tritium-labeled cholesterol assay. Taken together, our lipidomics analyses revealed that P116 can bind to different lipid species. While purified P116 mainly carried PGs, PEs and wax esters, the refilled P116 preferentially bound to PCs, SMs, and cholesterols. Notably, the composition of the lipid species in the refilled P116 was strikingly different than the serum lipid distribution. For example, highly abundant uncharged TGs did not bind to P116. Thus, P116 although displaying a large bandwidth of lipid uptake, it does show a preference for selective lipid species (**Figure 4c, 4d** and **Extended Data Table II**).

### P116 binds HDL between its N-terminal and core domains

Next, we performed cryoEM on a sample containing empty P116 and HDL. Of ∼46,000 particles that were identified as HDL, ∼25,000 were attached to P116. The resulting density at a resolution of 9 Å shows P116 interacting directly with HDL at the region between the N-terminal domain and the core (**Figure 4f**). In the reconstruction the density of P116 resembles the filled conformation, and the structure can be well fitted to the density map. Cryo-electron tomograms of whole *M. pneumoniae* cells indicate a similar arrangement of P116 with respect to the *Mycoplasma* membrane, although an unambiguous identification of the involved complexes is challenging due to the modest resolution (**Extended Data Figure 10**).

## Discussion

P116 is essential for the viability of the human pathogen *M. pneumoniae*^5^ and is the target of a strong antigenic response in infected patients^19^. The P116 structure has a previously unseen fold with a uniquely large hydrophobic cavity filled with ligands. Mass spectrometry and radioactivity transfer experiments confirm a lipid extraction from serum (FBS) and HDL. Further, the ligands are identified as essential lipids for the survival of the cells. Crosslinking mass spectrometry studies indicate one weak aminoacid-pair interaction between P116 and MPN161 (a protein of unknown function)^20^. Thus, while the involvement of other proteins in incorporating the extracted lipids into the *Mycoplasma* membrane cannot be excluded, it appears likely, given the observed conformational cycle upon lipid uptake, that P116 is also responsible for incorporation. Altogether, the P116 structure, along with our insights into different P116 conformations and the P116 complex formation with HDL, reveals a mechanism by which *Mycoplasmas* extract lipids from their environment and most likely incorporate them into their own membrane.

The transition from a full to an empty P116 molecule involves a ∼70% volume reduction of the hydrophobic cavity in concert with a wringing motion of the core domains. During this wringing motion, in which the monomers are each twisted in the opposite direction around their long axis, the hydrophobic cavities face almost opposite directions. Since, the N-terminal domain is anchored in the *Mycoplasma* membrane in vivo, the core domain is the one experiencing the high flexibility seen in our data sets. This enables an alternating motion of the core domain, in which each time one monomer of the core domain faces the *Mycoplasma* membrane (i.e. the one transferring lipids to the membrane) and the other monomer faces the environment (i.e. the one extracting lipids from the environment). This wringing motion can be repeated in a continuous manner. In this way, P116 could undergo a rolling movement on the *Mycoplasma* membrane, thus facilitating the transport of cholesterol and other essential lipids in an apparently simple and newly discovered way for lipid transporters.

*Mycoplasmas* have a minimal genome and are capable of incorporating many different lipids into their membrane^7,8^. The lipid-binding versatility shown by P116 enables a single molecular system to cope with the transport of diverse lipids required by *Mycoplasmas*. Although only *Mycoplasmas* share genes with similar sequences to P116, other microorganisms that require uptake of lipids from the environment, including clinically relevant bacterial species such as *Borrelia burgdorferi* and *Helicobacter pylori*^1^, may have similar -not yet discovered-systems to regulate their cholesterol homeostasis. Whether P116 shares functional similarities with other transfer proteins such as human cholesteryl ester transfer and phospholipid transfer proteins^21,22^ requires further investigation. However, the diversity and amount of lipids that P116 can bind appear to be unmatched by any other known prokaryotic or eukaryotic lipid carrier. This new understanding of bacterial lipid uptake opens possibilities for treatment of mycoplasma infections and may, for the first time^3^, allow the creation of a vaccine against *Mycoplasma pneumoniae*.

## Acknowledgements

We thank Laura Company and Irene Fernández-Vidal for their support during MALS and mass spectroscopy measurements, Antoni Iborra (Servei de Cultius Cel·lulars, Anticossos i Citometria, UAB) for his assistance with immunizing mice, and Rosa Pérez-Luque and David Aparicio for their constant support and discussions. We also thank the Central Electron Microscopy Facility, MPI of Biophysics in Frankfurt, which enabled us to collect the P116 empty dataset and in particular Sonja Welsch who assisted during the data collection. J.P. has been supported in part by grant BIO2017-84166-R from the Ministerio de Ciencia, Innovación y Universidades (Spain) to. I.F. was funded by Ministeriode Ciencia e Innovación (MICINN-Spain) grant PID2021-125632OB-C21. A.S.F. was supported by the Deutsche Forschungsgemeinschaft (FR 1653/14-1 for SM, FR 1653/6-3 for LS) and the Research Training Group iMOl (GRK 2566/1 for MPS).

## Author contributions

DV: Project initiation; preparation of the P116 gene synthesis; cloning of two constructs 13-957 and 30-657; expression and purification tests; adapting the best conditions for its stabilization. Expression, purification of mutant W681A and analysis by MALS. DV, LS, IF, ASF: Model building of the original, emptied and refilled P116 protein. DV and JMC: Emptying protocol. On-site assistance in the experimental preparation of the HDL-P116 interaction and the uptake of radioactive cholesterol (free and esterified). LS: Planned and carried out the single particle analysis; solved the P116 structure in the original, emptied and refilled state; proposed the mechanism of cholesterol uptake based on the structural data; solved the structure of P116 with HDL; model building of the empty P116 protein. SM: Planned and carried out the single particle sample preparation for the empty, refilled and the sample mixed with HDL. J.P. and M.M Obtaining hybridomas and monoclonal antibodies vs P116 protein. Immunolocation of P116 by epifluorescence microscopy, time-lapse microcinematography. MS: Advised on single particle experiments and structure determination procedures. JDL and JMC: Mass spectrometry analysis of all samples prepared for single particle analysis; a complete lipidomics analysis. IF: Designed and supervised research. ASF: Designed and supervised research. IF and ASF: Prepared the manuscript, with contributions from all authors.

## Data availability statement

Cryo-electron microscopy densities of the original P116 density map (3.3 Å), the empty P116 (4 Å) and the refilled P116 (3.5 Å) have been deposited in the EM Data Base under the accession codes XXXX, XXXX and XXXX, respectively. Model coordinates of empty and original P116 have been deposited in the PDB under the accession codes YYYY and YYYY, respectively.

## Author information

The authors declare no competing financial interests.

## Materials & Methods

### Bacterial strains, tissue cultures and growth conditions

*M. pneumoniae* M129 strain was grown in cell culture flasks containing SP4 medium and incubated at 37°C and 5% CO_2_. Surface-attached mycoplasmas were harvested using a cell scraper and resuspended in SP4 medium. To grow mycoplasma cells on IBIDI 8-well chamber slides, each well was seeded with about 10^5^ CFUs and incubated for 12–24 h in 200 μL SP4 supplemented with 3% gelatin.

NSI myeloma cells^23^ were grown in RPMI 1640 medium supplemented with 10% fetal bovine serum (FBS) and 50 μg mL^-1^ gentamycin (complete RPMI). Hybridomas were selected in complete RPMI supplemented with HAT media and BM-Condimed (Sigma Aldrich, St. Louis, USA).

### Cloning, expression, and purification of P116 constructs

Regions corresponding to the MPN213 gene from *M. pneumoniae* were amplified from synthetic clones (**Extended Data Table III**) using different primers for each construct: P116F_30_ and P116R_957_ for P116(30–957); P116F_13_ and P116R_957_ for P116(13–957); P116F_212_ and P116R_862_ for P116(212–862); and P116W_681_ to introduce mutation W681A (**Extended Data Table IV**). PCR fragments were cloned into the expression vector pOPINE (gift from Ray Owens; plasmid #26043, Addgene, Watertown, USA) to generate constructs, with a C-terminal His-tag. Recombinant proteins were obtained after expression at 22°C in B834 (DE3) cells (Merck, Darmstadt, Germany), upon induction with 0.6□mM IPTG at 0.8 OD600. Cells were harvested and lysed by French press in binding buffer (20 mM TRIS-HCl pH: 7.4, 40□mM imidazole and 150 mM NaCl) and centrifuged at 49,000□×□*g* at 4°C. Supernatant was loaded onto a HisTrap 5□ml column (GE Healthcare, Chicago, USA) that was pre-equilibrated in binding buffer and elution buffer (20 mM TRIS-HCl pH: 7.4, 400□mM imidazole and 150 mM NaCl). Soluble aliquots were pooled and loaded onto a Superdex 200 GL 10/300 column (GE Healthcare, Chicago, USA) in a protein buffer (20 mM TRIS-HCl□pH 7.4 and 150□mM NaCl).

To obtain empty P116, 2.6% Triton X-100 was added to the protein sample and incubated for 1.5 h at room temperature. Subsequent purification followed the same methodology described above, but also included a wash step with the binding buffer plus 1.3% Triton X-100, followed by extensive washing with at least 20 column volumes of wash buffer (20 mM TRIS-HCl pH: 7.4, 20□mM imidazole) before eluting the samples from the column. P116 was concentrated with Vivaspin 500 centrifugal concentrators (10,000 MWCO PES, Sartorius, Göttingen, Germany) to a final concentration of >0.5 mg/mL.

To refill P116 with lipids, the empty protein was incubated with approximately 1 ml FBS per mg P116 for 2 h at 30°C while still bound on the column. After extensive washing with at least 40 column volumes of wash buffer, elution and concentration were performed as described above.

### HDL isolation and determination of cholesterol transfer rate

Human HDL (density 1.063–1.210 g/mL) was isolated from plasma of healthy donors via sequential gradient density ultracentrifugation, using potassium bromide for density adjustment, at 100,000 *g* for 24 h with an analytical fixed-angle rotor (50.3, Beckman Coulter, Fullerton, CA, USA). The amount of cholesterol and apolipoprotein A1 were determined enzymatically and by an immunoturbidimetric assay, respectively, using commercial kits adapted for a COBAS 6000 autoanalyzer (Roche Diagnostics, Rotkreuz, Switzerland). Radiolabeled HDLs were prepared as previously described^24^. Briefly, 10 μCi of either [1,2-^3^H(N)] free cholesterol or [1,2-^3^H(N)]cholesteryl oleate (Perkin Elmer, Boston, MA) were mixed with absolute ethanol, and the solvent was dried under a stream of N_2_. HDL (0.5 mL, 2.25 g/L of ApoA1) was added to the tubes containing the radiotracers, as appropriate, and then incubated for 16 h in a 37°C bath. The labeled HDLs (both ^3^H-cholesterol-containing and ^3^H-cholesteryl oleate-containing HDLs) were re-isolated by gradient density ultracentrifugation at 1.063–1.210 g/mL and dialyzed against PBS via gel filtration chromatography. Specific activities of ^3^H-cholesterol-containing and ^3^H-cholesteryl oleate-containing HDLs were 1221 and 185 counts per minute (cpm)/nmol, respectively. The cholesterol transfer to P116 (1 g/L) was measured after adding either [^3^H] free cholesterol-containing or [^3^H]cholesteryl oleate-containing HDL (0.5 g/L of APOA1) and incubating for 2 h at 37°C. HDL and P116 were separated by a HisTrap HP affinity column. The radioactivity associated with each P116 and HDL fraction was measured via liquid scintillation counting. The percentage of [^3^H]cholesterol transferred per mL was determined for each condition. The specific activities for each radiotracer were used to calculate the amount of free cholesterol and cholesteryl ester transferred from HDL to P116.

### Size exclusion chromatography and multi-angle light scattering (SEC-MALS)

Molecular weights were measured from P116 samples using a Superose 6 10/300 GL (GE Healthcare, Chicago, USA) column in a Prominence liquid chromatography system (Shimadzu, Kyoto, Japan) connected to a DAWN HELEOS II multi-angle light scattering (MALS) detector and an Optilab T-REX refractive index (dRI) detector (Wyatt Technology, Santa Barbara, USA). ASTRA 7 software (Wyatt Technology) was used for data processing and analysis. An increment of the specific refractive index in relation to concentration changes (dn/dc) of 0.185 mL/g (typical of proteins) was assumed for calculations.

### Matrix-assisted laser desorption/ionization-mass spectrometry (MALDI-TOF)

All samples were mixed in a 1:1 ratio with either DHB or sDHB (Bruker Daltonics, Germany) matrix solution (50 mg·ml^-1^ in 50% Acetonitrile (ACN), 50% water and 0.1% TFA). Subsequently 1 μl aliquots of the mixture were deposited on a BigAnchor MALDI target (Bruker Daltonics, Germany) and allowed to dry and crystallize at ambient conditions. Unless stated otherwise, all reagents and solvents were obtained from Sigma Aldrich, Germany.

MS spectra were acquired on a rapifleX MALDI-TOF/TOF (Bruker Daltonics, Germany) in the mass range from 20.000-120.000 m/z in linear positive mode and in the mass range from 100-1600 m/z in reflector positive mode. The Compass 2.0 (Bruker, Germany) software suite was used for spectra acquisition and processing.

### Lipidomics analysis (LC-TIMS-MS/MS)

Samples were extracted using a modified MTBE/Methanol extraction protocol, and submitted to LC-nanoESI-IMS-MS/MS analysis using a Bruker NanoElute UHPLC coupled to a Bruker TimsTOF Pro 2 mass spectrometer operated in DDA-PASEF mode. In brief, 40 min gradients on PepSep C18 columns (1.9A, 75µm ID, 15cm length) were recorded in positive and negative ion mode. Data were analysed using the MS-DIAL pipeline (version 4.9).

### Single-particle cryoEM

For single-particle cryoEM, a 3.5 µl drop of purified P116 (100–400 µg/mL in 20 mM Tris, pH 7.4 buffer or 600 µg/mL in 20 mM Tris, 2 mM CHAPSO, pH 7.4 buffer) or P116 mixed with HDL (250 µg/mL P116 and 1116 µg/mL HDL in 20 mM Tris, pH 7.4 buffer) was applied to a (45 s) glow-discharged R1.2/1.3 C-flat grid (Electron Microscopy Science, Hatfield, USA), and plunge-frozen in liquid ethane (Vitrobot Mark IV, Thermo Scientific, Waltham, USA) at 100% relative humidity, 4 °C, nominal blot force –3, wait time 45 s, with a blotting time of 12 s. Before freezing, Whatman 595 filter papers were incubated for 1 h in the Vitrobot chamber at 100% relative humidity and 4°C.

Dose-fractionated Movies of P116, P116 refilled and P116 mixed with HDL were collected with SerialEM v3.8^25^ at a nominal magnification of 130,000x (1.05 Å per pixel) in nanoprobe EFTEM mode at 300 kV with a Titan Krios (Thermo Scientific, Waltham, USA) electron microscope equipped with a GIF Quantum S.E. post-column energy filter in zero loss peak mode and a K2 Summit detector (Gatan Inc., Pleasanton, USA). For P116, P116 refilled and P116 with HDL a total of 4376, 4019 and 3114 micrographs with 34, 29 and 30 frames per micrograph and a frame time of 0.2 s were collected. The camera was operated in dose-fractionation counting mode with a dose rate of ∼8 electrons per Å ^2^ s^-1^, resulting in a total dose of 50 electrons per Å^2^ s^-1^. Defocus values ranged from –1 to –3.5 µm.

For P116 empty, dose-fractionated Movies were collected using EPU 2.12 (Thermo Scientific, Waltham, USA) at a nominal magnification of 105,000x (0.831 Å per pixel) in nanoprobe EFTEM mode at 300 kV with a Titan Krios G2 electron microscope (Thermo Scientific, Waltham, USA), equipped with a BioQuantum-K3 imaging filter (Gatan Inc., Pleasanton, USA), operated in zero loss peak mode with 20 eV energy slit width. In total 15,299 micrographs with 50 frames per micrograph and frame time of 0.052 s were collected. The K3 camera was operated in counting mode with a dose rate of ∼ 16 electrons per A^2^ s^-1^, resulting in a total dose of 50 electrons per Å^2^ s^-1^. Defocus values ranged from -0.8 to -3.5 µm.

CryoSPARC v3.2^26^ was used to process the cryoEM data, unless stated otherwise. Beam-induced motion correction and CTF estimation were performed using CryoSPARC’s own implementation. Particles were initially clicked with the Blob picker using a particle diameter of 200–300 Å. Particles were then subjected to unsupervised 2D classification. For the final processing, the generated 2D averages were taken as templates for the automated particle picking, for the processing of P116 with HDL no template picking was performed. In total, 3,463,490, 4,532,601 particles, 2,930,863 particles and 262,981 particles were picked and extracted with a binned box size of 256 pixels for P116, P116 empty, P116 refilled and P116 with HDL respectively. False-positive picks were removed by two rounds of unsupervised 2D classification. The remaining 1,324,330 particles (P116), 1,140,275 particles (P116 empty), 1,311,526 particles (P116 refilled) and 46,277 particles (P116 with HDL) were used to generate an ab initio reconstruction with three classes followed by a subsequent heterogeneous refinement with three classes. For the final processing, 1,315,362 particles (P116), 633,332 particles (P116 empty), 1,311,526 particles (P116 refilled) and 46,277 particles (P116 with HDL) were used. For the remaining particles, the beam-induced specimen movement was corrected locally.

The CTF was refined per group on the fly within the non-uniform refinement. The obtained global resolution of the homodimer was 3.3 Å (P116), 4 Å (P116 empty), and 3.5 Å (P116 refilled) (**Extended Data Figure 1 and Extended Data Table I**). To analyze the sample in regard to its flexibility the particles were subjected to the 3D variability analysis of cryoSPARC which was used to display the continuous movements of the protein.

### Cryo-electron tomography of *M. pneumoniae*

*M. pneumoniae* M129 cells of an adherently growing culture were scraped in a final volume of 1 ml of SP4 medium and washed three times in PBS. This solution was mixed with fiducial markers (Protein A conjugated to 5 nm colloidal gold: Cell biology department, University Medical Center Utrecht, The Netherlands). From this stock a 3.5 µl drop was applied to a (45 s) glow-discharged R1.2/1.3 C-flat grid (Electron Microscopy Science, Hatfield, USA), and plunge-frozen in liquid ethane (Vitrobot Mark IV, Thermo Scientific, Waltham, USA) at 100% relative humidity, 4 °C, nominal blot force –1, with a blotting time of 10 s.

Tilt-series were recorded using SerialEM v3.8^25^ at a nominal magnification of 105,000x (1.3 Å per pixel) in nanoprobe EFTEM mode at 300 kV with a Titan Krios (Thermo Scientific, Waltham, USA) electron microscope equipped with a GIF Quantum S.E. post-column energy filter in zero loss peak mode and a K2 Summit detector (Gatan Inc., Pleasanton, USA). The total dose per tomogram was 120 e^-^/ Å^2^, the tilt series covered an angular range from -60° to 60° with an angular increment of 3° and a defocus set at -3 µm. Tomograms were reconstructed by super-sampling SART^27^ with a 3D CTF correction^28^.

### P116 model building and refinement

The initial tracing of the core domain was performed manually with Coot^29^. It contained numerous gaps and ambiguities that were slowly polished by alternating cycles of refinement using the “Real Space” protocol in the program Phenix^30,31^ and manual reinterpretation and rebuilding with Coot. The tracing and assignment of specific residues in the N-terminal domain were very difficult due to the low local resolution of the map for this domain, and only a partial interpretation was achieved. Using Robetta and AlphaFold ^14^ we obtained different predictions of the N-terminal domain structure using different parts of the sequence. The highest ranked predictions, selected using the partial experimental structure already available, were obtained with AlphaFold for residues 81–245, which allowed us to complete the building of the N-terminal domain according to the cryoEM map. The RMS deviation between the AlphaFold prediction and the experimental model was 2.6 Å for 104 (63%) structurally equivalent residues. Some residues at the N-end of the N-terminal domain were difficult to identify and were represented as alanines in the final model. The whole P116 model was then refined using Phenix, and the final refined structure was deposited in the EMDB with code XXXX **(Extended Data Table II)**.

### Polyclonal and monoclonal antibody generation

Two BALB/C mice were serially immunized with four intraperitoneal injections, each one containing 150 μg of recombinant P116 ectodomain (residues 30–957) in 200 μL of PBS with no adjuvants. The last injection was delivered four days before splenectomy. Isolated B lymphocytes from the immunized mice were fused to NSI myeloma cells^23^ to obtain stable hybridoma cell lines producing monoclonal antibodies, as previously described^32^. Supernatants from hybridoma cell lines derived from single fused cells were first investigated by indirect ELISA screening against the recombinant P116 ectodomain. Positive clones were also tested by Western blot against protein profiles from *M. pneumoniae* cell lysates and by immunofluorescence using whole, non-permeabilized M. pneumoniae cells (see below). Only those clones with supernatants revealing a single 116 kDa band in protein profiles and also exhibiting a consistent fluorescent staining of *M. pneumoniae* cells were selected and used in this work. Polyclonal sera were obtained by cardiac puncture of properly euthanized mice just before splenectomy and titred using serial dilutions of the antigen. The titre of each polyclonal serum was determined as the IC_50_ value from four parameter logistic plots and found to be approximately 1/4000 for both sera. Polyclonal anti-P1 and anti-P90/P40 antibodies were obtained by immunizing two BALB/C mice with recombinant P1 and P90/P40 proteins^33^, respectively, as described above. The titres obtained for polyclonal anti-P1 antibodies and anti-P90/P40 antibodies were approximately 1/2500 and 1/3000, respectively.

### Immunofluorescence microscopy

The immunofluorescence staining of mycoplasma cells on chamber slides was similar to previously described^34^, with several modifications. Cells were washed with PBS containing 0.02% Tween 20 (PBS-T) prewarmed at 37°C, and each well was fixed with 200 μL of 3% paraformaldehyde (wt/vol) and 0.1% glutaraldehyde. Cells were washed three times with PBS-T, and slides were immediately treated with 3% BSA in PBS-T (blocking solution) for 30 min. The blocking solution was removed, and each well was incubated for 1 h with 100 μL of the primary antibodies diluted in blocking solution. For P116 polyclonal sera, we used a 1/2000 dilution; a 1/10 dilution was used for monoclonal antibodies from hybridoma supernatants. Wells were washed three times with PBS-T and incubated for 1 h with a 1/2000 dilution of a goat anti-mouse Alexa 555 secondary antibody (Invitrogen, Waltham, USA) in blocking solution. Wells were then washed three times with PBS-T and incubated for 20 min with 100 μL of a solution of Hoechst 33342 10 μg/μL in PBS-T. Wells were finally washed once with PBS-T and replenished with 100 μL of PBS before microscopic examination. Cells were observed by phase contrast and epifluorescence in an Eclipse TE 2000-E inverted microscope (Nikon, Tokyo, Japan). Phase contrast images, 4’,6-diamidino-2-phenylindole (DAPI, excitation 387/11 nm, emission 447/60 nm) and Texas Red (excitation 560/20 nm, emission 593/40 nm) epifluorescence images were captured with an Orca Fusion camera (Hamamatsu, Hamamatsu, Japan) controlled by NIS-Elements BR software (Nikon, Tokyo, Japan).

### Time-lapse microcinematography

The effect of anti-P116 antibodies and anti-P1 polyclonal serum on mycoplasma cell adhesion was investigated by time-lapse cinematography of *M. pneumoniae* cells growing on IBIDI 8-well chamber slides. Before observation, medium was replaced with PBS containing 10% FBS and 3% gelatin prewarmed at 37°C. A similar medium has been used to test the effect of P1 antibodies on mycoplasma adhesion and gliding motility^35^. After incubation for 10 min at 37°C and 5% CO_2_, the slide was placed in a Nikon Eclipse TE 2000-E inverted microscope equipped with a Microscope Cage Incubation System (Okolab, Pozzuoli, Italy) at 37°C. Images were captured at 0.5 s intervals for a total observation time of 10 min. After the first 60 s of observation, the different antibodies were dispensed directly into the wells. The frequencies of motile cells and detached cells before the addition of antibodies were calculated from the images collected between 0 and 60 s of observation. The frequencies of motile cells and detached cells after the addition of antibodies were calculated from the images collected in the last minute of observation.

