## Extended Data for "The immunodominant protein P116 extracts cholesterol and other essential lipids"

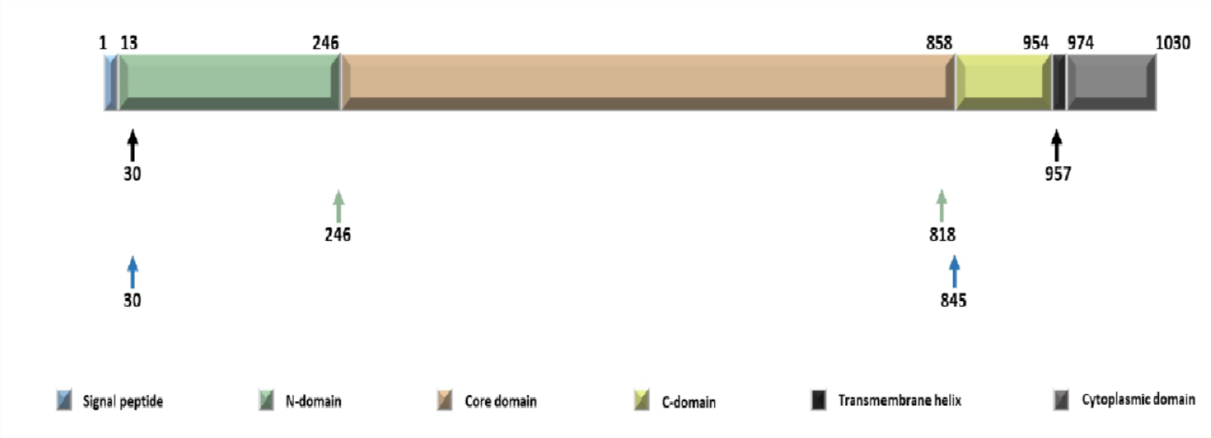

**Extended Data Figure 1: P116 constructs.**

Overview of the different constructs (30 - 957, 30 – 818, 246 - 845) used for expression. For expression purposes a His-tag (KHHHHH) was added at the C-terminus. For the structural analysis by cryoEM the construct from 30-957 was used.

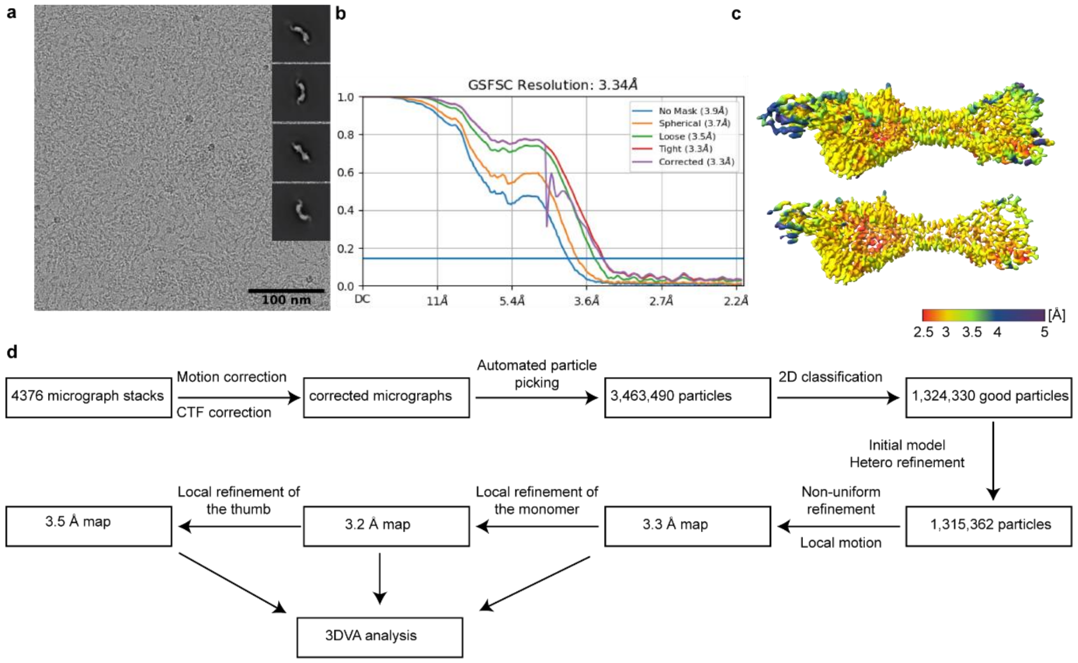

**Extended Data Figure 2: Overview of cryoEM processing of P116. (a)** Representative Micrograph and obtained 2D classes used for template picking. **(b)** Fourier shell correlation of P116 reporting a final resolution of 3.3 Å according to the 0.143 cut-off criteria. **(c)** Local resolution map of the cryoEM denisty map of P116 ranging from 2.5 to 5 Å. **(d)** Schematic processing overview for the P116 construct. The processing for P116 empty, P116 refilled and P116 + HDL was carried out in a similar manner.

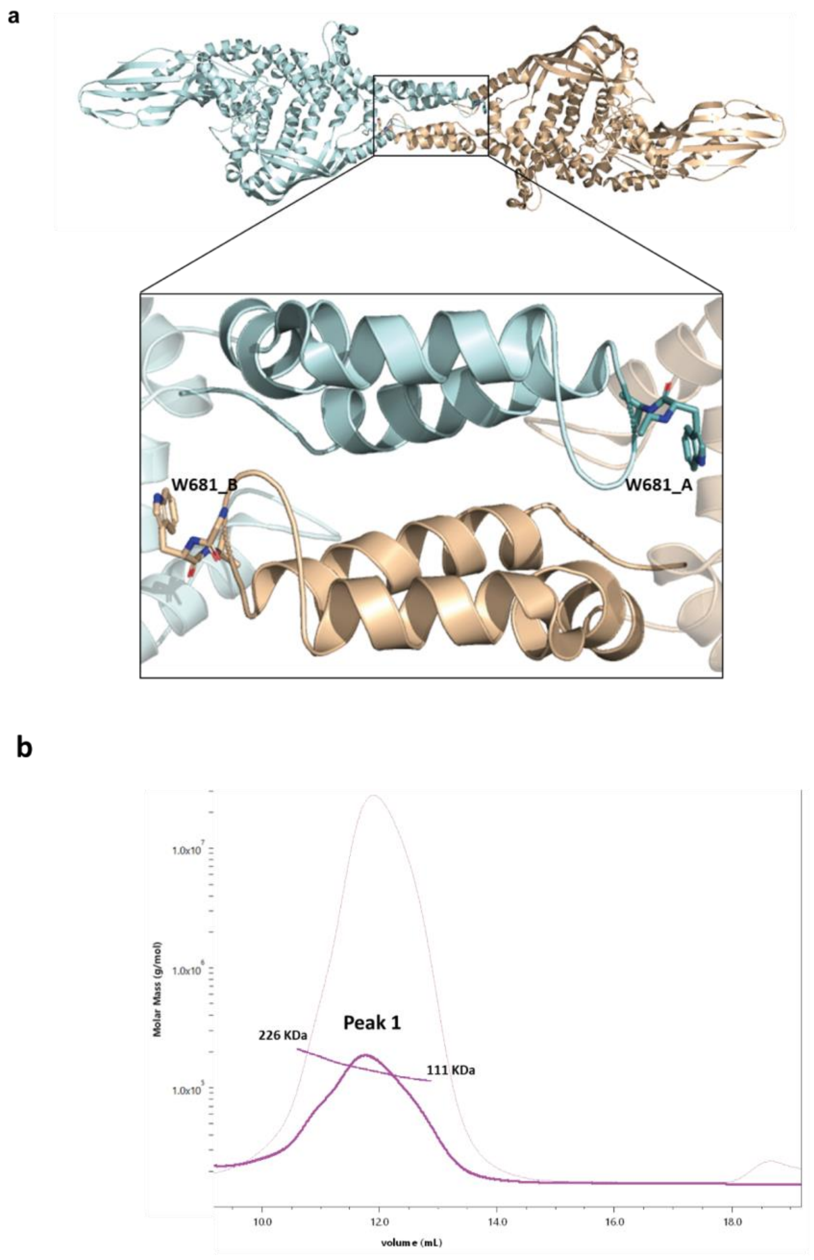

| Sample | Mw (KDa) | Polydispersity (Mw/Mn) | Mass fraction (%) |
|---|---|---|---|
| Peak 1 | 143,4 +/- 0,27 | 1,02 +/- 0,00 | 98,8 |

**Extended Data Figure 3: Dimerization interface of P116. (a)** Detail of the dimerization zone with the two monomers and the TRP-681 residue of both chains contacting the opposite one. **(b)** SEC-MALS of P116 W681A. The greatest polydispersity of this sample is observed, which oscillates between a very large MW (molecular weight) range, and the clear decrease in size with respect to the WT due to the now predominance of the monomeric state.

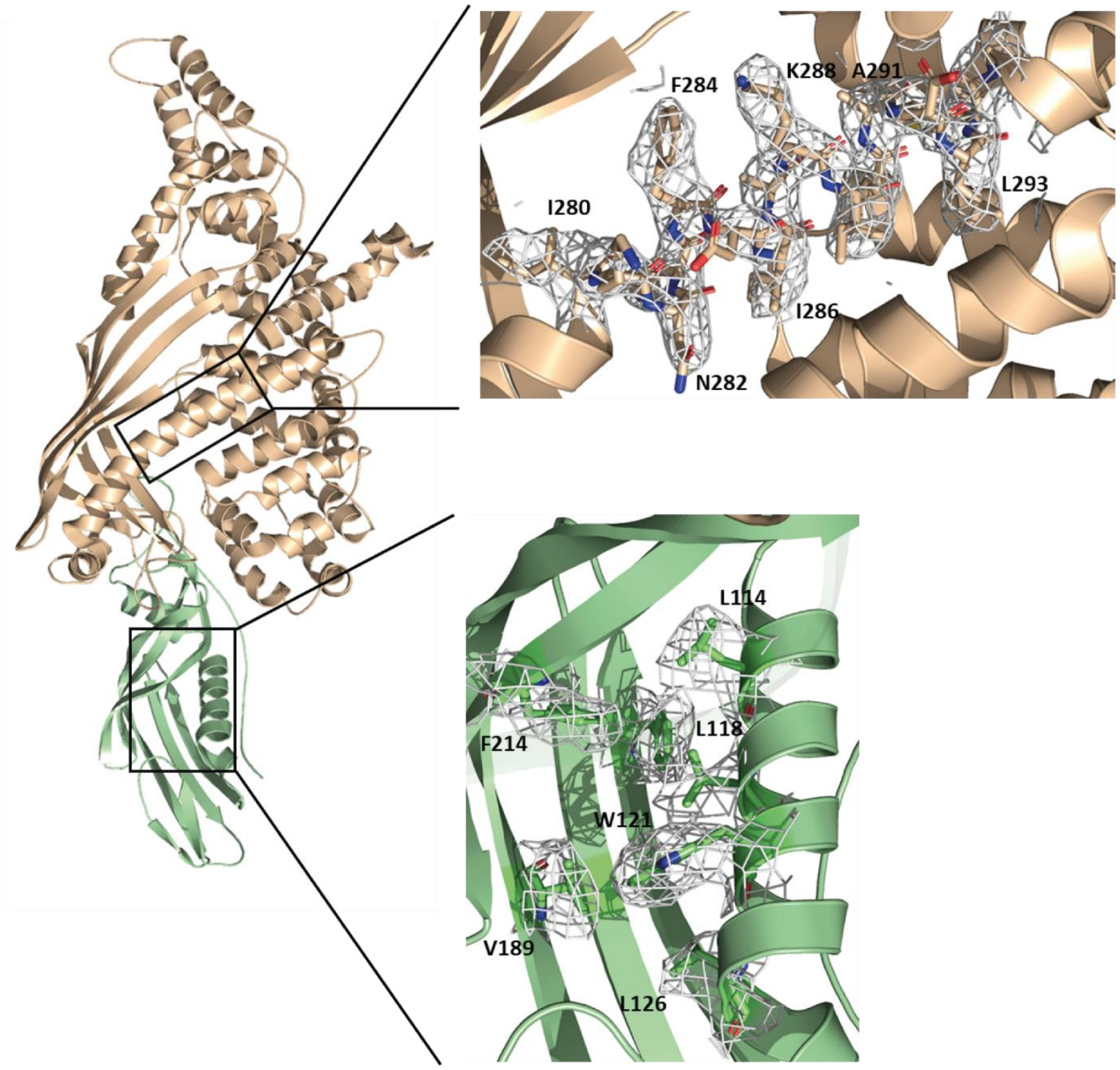

**Extended Data Figure 4: Model building of P116.** Exemplary superimpositions of the cryoEM density and the PDB model for the core and N-terminal domain.

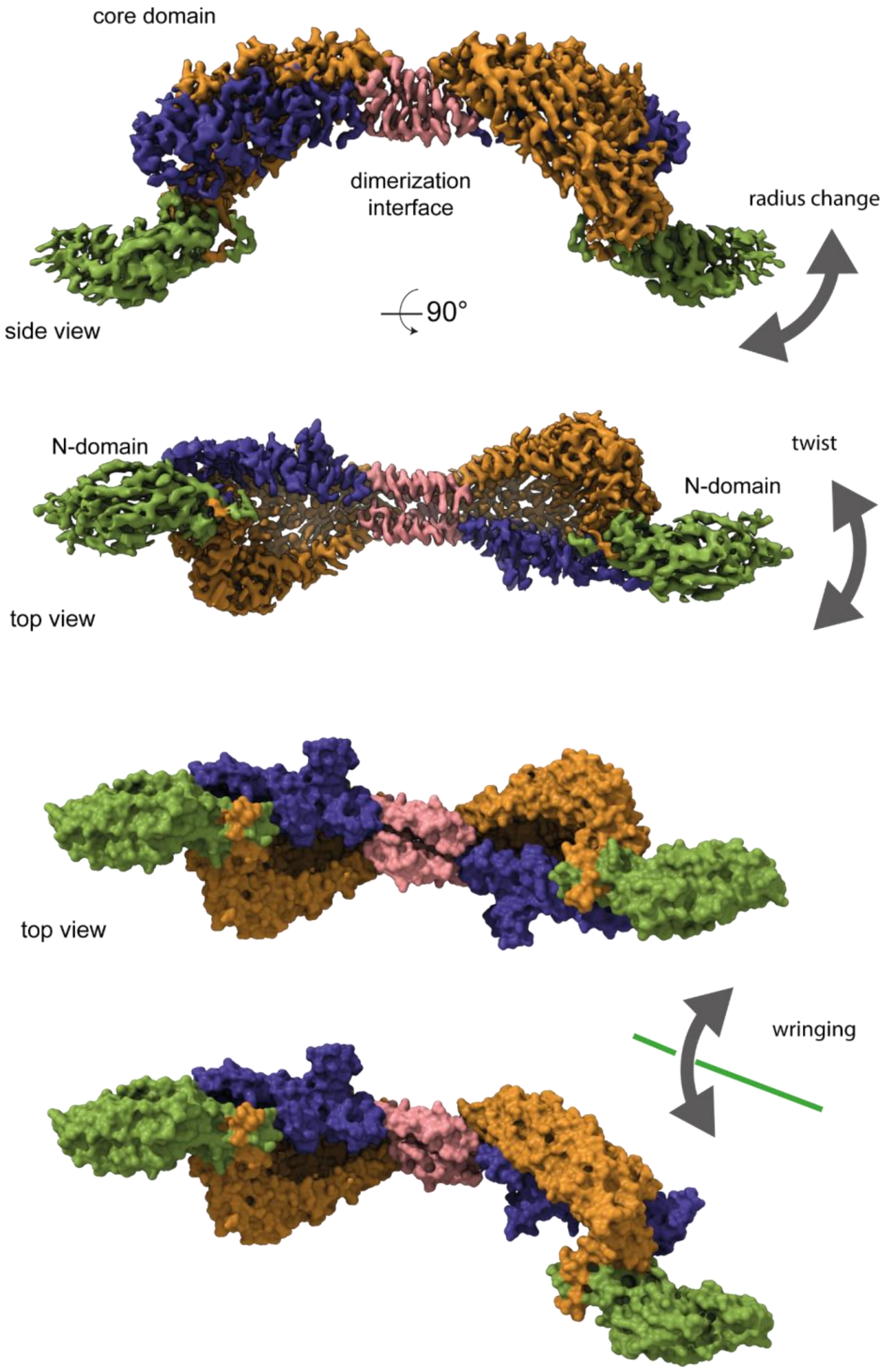

**Extended Data Figure 5: Flexibility of P116.** CryoEM classes display different flexibility modes.

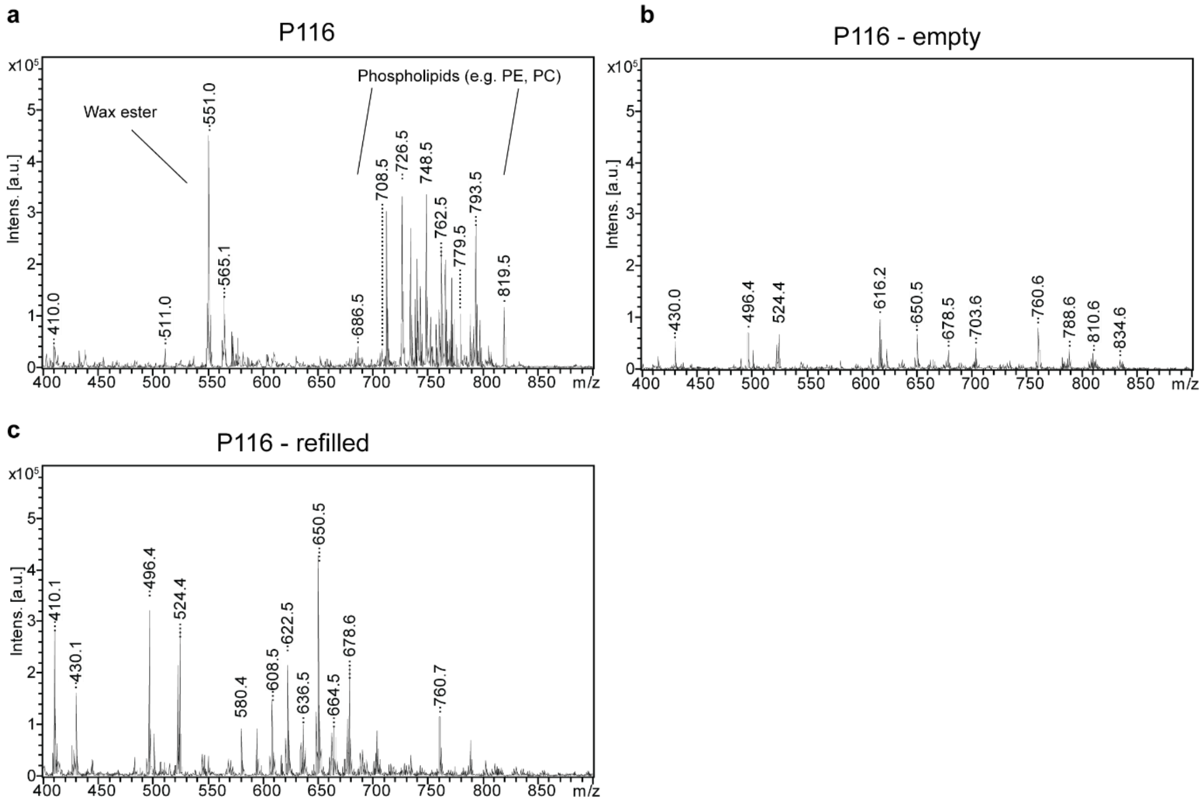

**Extended Data Figure 6: MALDI-TOF of P116, P116 empty and P116 refilled (a-c)**
Individual low molecular weight MALDI-MS spectra of P116, P116-empty and P116-refilled as displayed in in figure 4D.

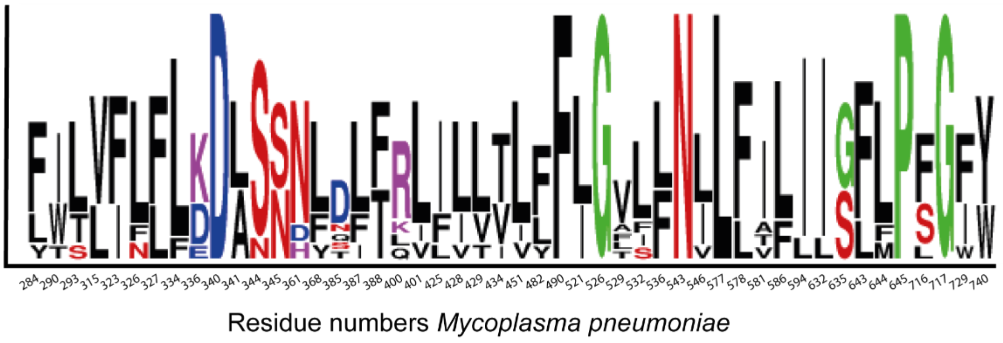

**Extended Data Figure 7: Conserved (minimum 55 %) amino acids pointing inwards the hydrophobic cavity.** Black indicates amino acids with a hydrophobic character. Red indicates amino acids with a hydrophilic character. Blue indicates amino acids with a negative side chain. Purple indicates amino acids with a positive side chain. Green indicates other amino acids.

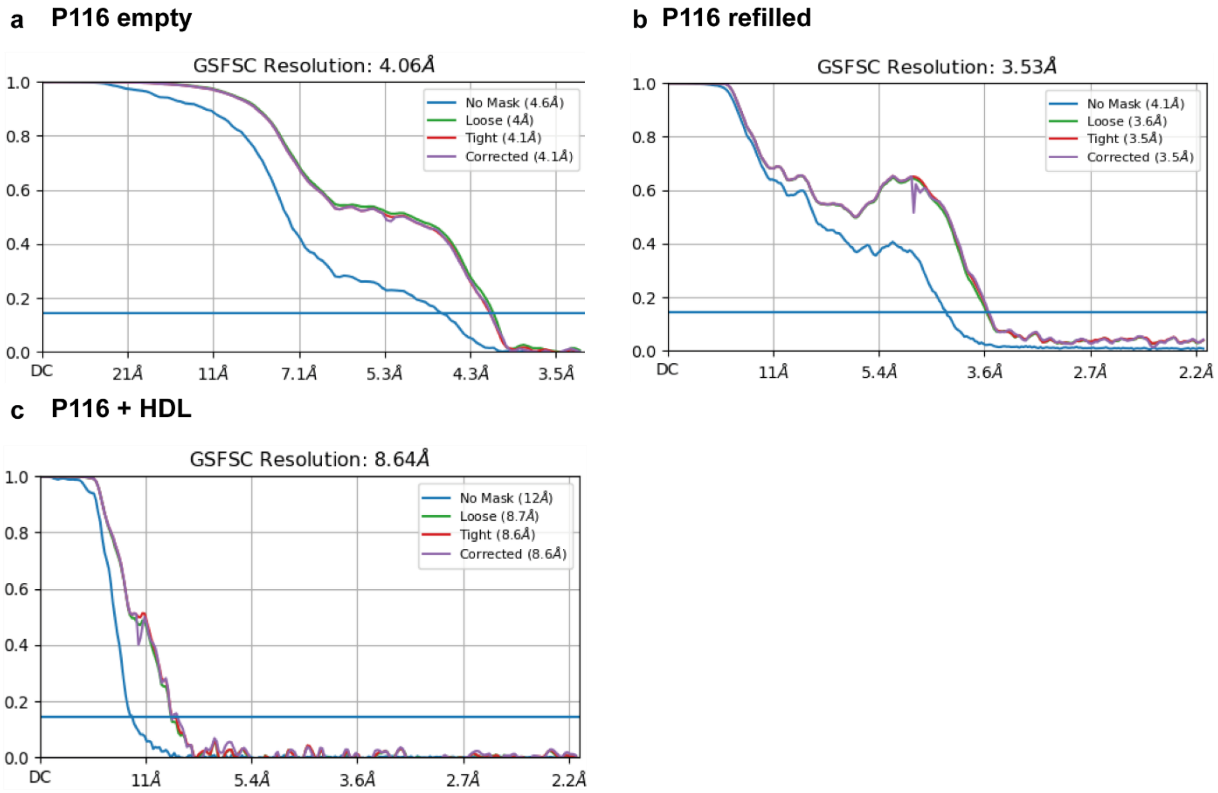

**Extended Data Figure 8: Fourier shell correlations of P116 empty, P116 refilled and P116 + HDL. (a)** Final reported resolution of P116 empty at 4 Å according to the 0.143 cut-off criteria. **(b)** Final reported resolution of P116 refilled at 3.5 Å according to the 0.143 cut-off criteria. **(c)** Final reported resolution of P116 + HDL at 9 Å according to the 0.143 cut-off criteria.

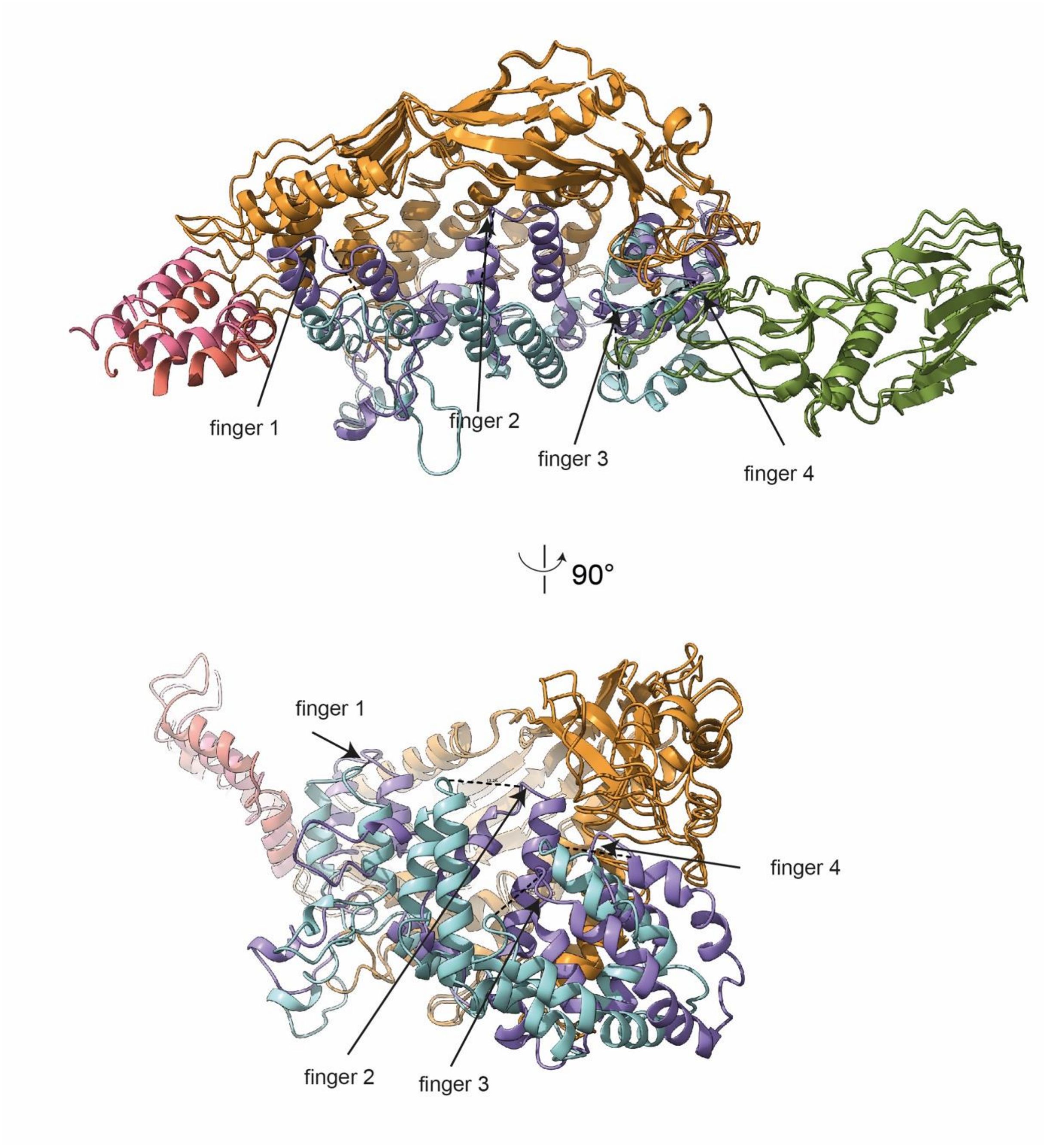

**Extended Data Figure 9: Conformational change between P116 and P116 empty.** Superimposition of one subunit of P116 with one subunit of P116 empty. The movement of the fingers 1-4 is indicated by dashed lines. Side view of the ribbon representation of the empty and full P116 shows that the fingers (in purple) have come closer to the core domain, massively reducing the available volume. Their new position is markedly different compared to the full P116 (shown in light blue). Finger 1 moved 8Å sideways and towards the core, finger 2 has moved 13 Å towards the core and Finger 3 has moved 12 Å towards the core.

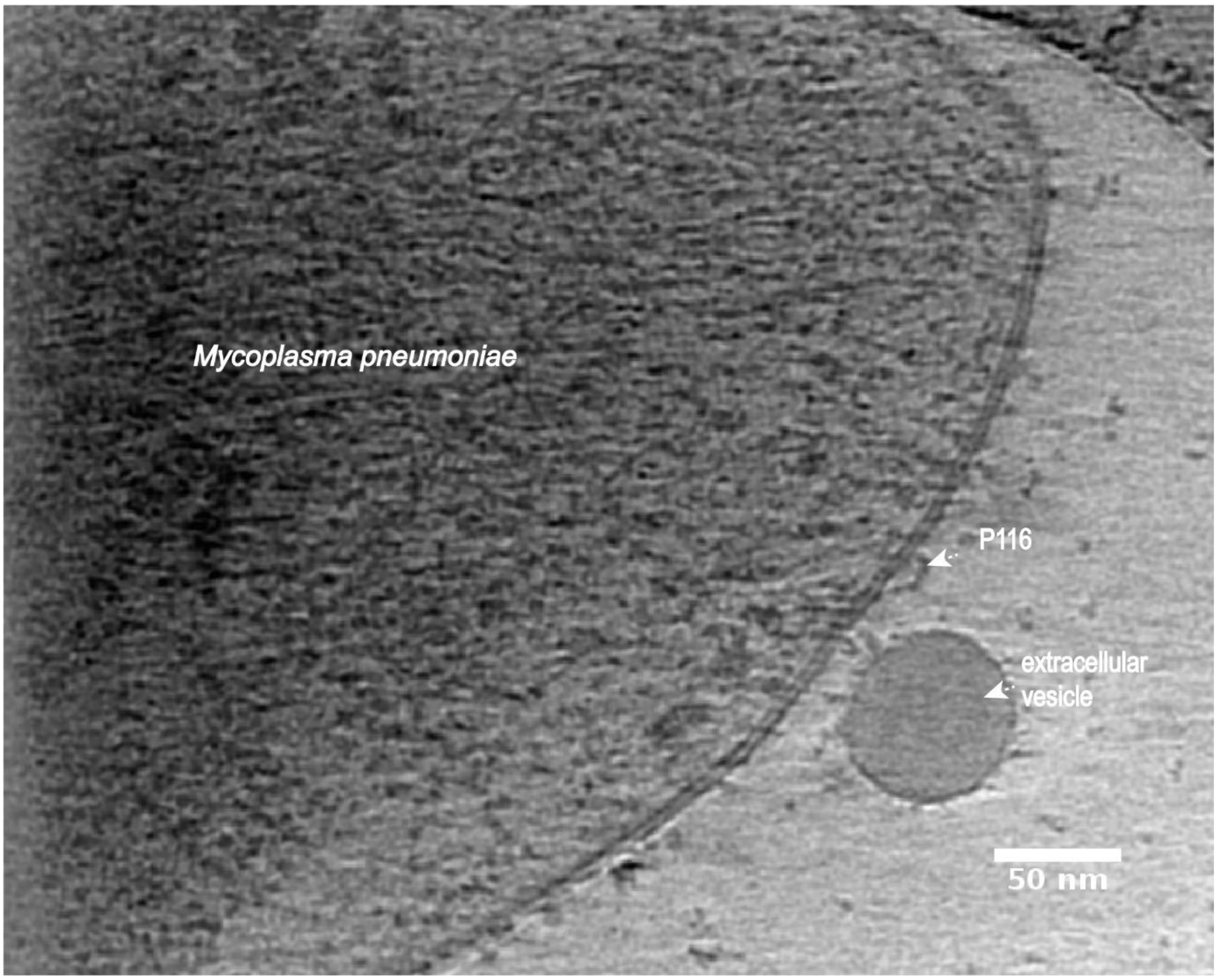

**Extended Data Figure 10: Cryo-electron tomogram of a *M. pneumoniae* cell.** Ortho slice of a tomographic reconstruction of a *M. pneumoniae* cell depicting a possible P116 protein on the surface of the *M. pneumoniae* membrane. Tomogram was acquired at a defocus of -3 µm.

**Extended Data Table I. CryoEM data collection, refinement and validation statistics**

| | P116<br>EMDB | P116 empty<br>EMDB | P116 refilled<br>EMDB | P116 + HDL<br>EMDB |
|---|---|---|---|---|
| **<u>Data collection</u>** | | | | |
| **Microscope** | FEI Titan Krios | FEI Titan Krios | FEI Titan Krios | FEI Titan Krios |
| **Detector** | Gatan K2 Summit | Gatan K3 Summit | Gatan K2 Summit | Gatan K2 Summit |
| **Acquistion Software** | SerialEM 3.8 | EPU 2.12 | SerialEM 3.8 | SerialEM 3.8 |
| **Magnification** | 130,000x | 105,000x | 130,000x | 130,000x |
| **Pixel size (Å)** | 1,05 | 0,831 | 1,05 | 1,05 |
| **Total electron dose ($e^-$/$Å^2$)** | 50 | 50 | 50 | 50 |
| **Dose rate ($Å^2$/$s^{-1}$)** | 8,2 | 8,2 | 8,2 | 30 |
| **Number of frames** | 34 | 50 | 29 | -1 to -3.5 |
| **Defocus range (µm)** | -1 to -3.5 | -1 to -3.5 | -1 to -3.5 | 3114 |
| **Micrographs used** | 4367 | 15299 | 4019 | |
| **<u>Processing</u>** | | | | |
| **Software** | cryoSPARC v3.3.2 | cryoSPARC v3.3.2 | cryoSPARC v3.3.2 | cryoSPARC v3.3.2 |
| **Motion correction** | cryoSPARC v3.3.2 | cryoSPARC v3.3.2 | cryoSPARC v3.3.2 | cryoSPARC v3.3.2 |
| **CTF estimation** | cryoSPARC v3.3.2 | cryoSPARC v3.3.2 | cryoSPARC v3.3.2 | cryoSPARC v3.3.2 |
| **Total extracted particles** | 3.463.490 | 4.532.601 | 2.930.863 | 262.981 |
| **After 2D classification** | 1.324.330 | 1.140.275 | 1.311.526 | 46.277 |
| **Number of refined particles** | 1.315.362 | 633.322 | 1.311.526 | 46.277 |
| **Symmetry** | / | / | / | / |
| **Map sharpening B factor** | -117 | -105 | -131 | -490 |
| **Resolution (Å) FSC 0.143** | 3,3 | 4 | 3,5 | 8.6 |

| **Refinement** | **P116** |
|---|---|
| **Initial model used (PDB code)** | / |
| **Model resolution (Å)** | 3.2 |
| **FSC threshold** | 0.143 |
| **Model resolution range (Å)** | 2.5-5.0 |
| **Map sharpening *B* factor ($Å^2$)** | -94 |
| **Model composition** | |
| **Non-hydrogen atoms** | 12772 |

| | |
|---|---|
| **Protein residues** | 1618 |
| **Ligands** | 0 |
| ***B* factors (Å$^2$)** | |
| **Protein** | 51.22 |
| **Ligand** | 0 |
| **R.m.s. deviations** | |
| **Bond lengths (Å)** | 0.003 |
| **Bond angles (°)** | 0.651 |
| **Validation** | |
| **MolProbity score** | 2.18 |
| **Clashscore** | 12 |
| **Poor rotamers (%)** | 0.14 |
| **Ramachandran plot** | |
| **Favored (%)** | 89.34 |
| **Allowed (%)** | 10.53 |
| **Disallowed (%)** | 0.13 |

**Extended Data Table II: Identified lipid compounds from of P116, P116-empty, P116-refilled and serum in positive and negative mode as used for heatmap generation.**

| Lipids positive mode | | | | | Lipids negative mode | | | | |
|---|---|---|---|---|---|---|---|---|---|
| Lipid ID | **Intensity P116** | **Intensity P116 - empty** | **Intensity P116 - refilled** | **Intensity serum** | **Lipid ID** | **Intensity P116** | **Intensity P116 - empty** | **Intensity P116 - refilled** | **Intensity serum** |
| AHexCer 41:0;3O | 10297 | 16374 | 13072 | 4881 | **ASG 29:1;O; Hex;FA 18:2** | 0 | 172 | 222 | 233 |
| AHexCer 42:2;3O | 0 | 303 | 29554 | 14318 | **Cer 34:1;3O** | 1980 | 6952 | 6374 | 2812 |
| AHexCer 43:6;3O | 68476 | 31 | 216 | 49285 | **Cer 42:0;4O** | 7028 | 7183 | 6499 | 2567 |
| AHexCer 45:6;3O | 129 | 3957 | 25 | 118 | **Cer 40:1;4O** | 945 | 984 | 2197 | 560 |
| AHexCer 50:11;3O | 3846 | 1697 | 4050 | 1781 | **Cer 41:0;3O** | 5083 | 5119 | 2345 | 1209 |
| ASG 29:2;O; Hex;FA 21:4 | 39904 | 4293 | 4810 | 1384 | **DGTS 39:10** | 37 | 33 | 1590 | 1194 |
| BMP 45:8 | 1558 | 15978 | 4520 | 271 | **LPE O-18:1** | 20 | 65 | 1247 | 3709 |
| SE 28:2/26:5 | 35039 | 13117 | 15374 | 3614 | **LPE O-20:1** | 35 | 71 | 7686 | 6318 |
| CE 20:3 | 16 | 758 | 12406 | 18956 | **LPE O-22:1** | 2 | 22 | 4260 | 4925 |
| CE 22:6 | 15 | 3241 | 26722 | 34996 | **LPG O-9:0** | 597 | 9668 | 8151 | 14248 |
| Cer 36:6;4O | 47440 | 103205 | 39840 | 44460 | **PC O-34:3;1O** | 24 | 207 | 10427 | 1885 |
| Cer 49:0;4O | 1685 | 2675 | 2395 | 1170 | **PC O-32:1;3O** | 11 | 12 | 49322 | 14761 |

| | | | | | | | | | |
|---|---|---|---|---|---|---|---|---|---|
| Cer 49:10;4O | 4360276 | 1364375 | 876 | 12937 | **PC O-32:0;3O** | 5 | 37 | 3451 | 958 |
| Cer 42:0;3O | 30798 | 30758 | 18380 | 8938 | **PC O-34:3;3O** | 119 | 10 | 5534 | 21113 |
| Cer 52:6;3O | 31238 | 19480 | 36506 | 67177 | **PC O-34:2;3O** | 14 | 39 | 53835 | 19019 |
| Cer 38:7;3O | 141116 | 125810 | 300 | 84689 | **PC O-34:1;3O** | 0 | 18 | 9960 | 2987 |
| Cer 38:1;3O | 11731 | 13247 | 14750 | 55589 | **PC O-37:5;1O** | 36 | 182 | 24629 | 10958 |
| Cer 52:7;3O | 31117 | 20209 | 33117 | 68691 | **PC O-38:5;1O** | 30 | 53 | 47834 | 2761 |
| Cer 35:6;2O | 71957 | 89844 | 53769 | 53725 | **PC O-36:4;3O** | 33 | 13 | 9183 | 7297 |
| Cer 34:8;2O | 221480 | 137961 | 278201 | 74577 | **PC O-36:3;3O** | 34 | 23 | 3383 | 1306 |
| Cer 34:8;2O | 39303 | 502 | 219635 | 1113 | **PC O-36:3;3O** | 6 | 11 | 14849 | 4665 |
| Cer 34:8;2O | 346597 | 242834 | 234 | 302185 | **PC O-32:0** | 6 | 6 | 7082 | 8969 |
| Cer 44:4;2O | 5421 | 4110 | 18587 | 30994 | **PC O-36:3** | 3 | 10 | 669 | 84 |
| CoQ6 | 12299 | 276 | 8354 | 169 | **PC O-38:7** | 49 | 67 | 3119 | 1107 |
| DG 26:5 | 166408 | 50223 | 233 | 86 | **PC O-38:5** | 0 | 29 | 3468 | 5014 |
| DG 28:2 | 452685 | 403343 | 481239 | 324957 | **PC O-42:6** | 11 | 2 | 2376 | 2904 |
| DG 28:0 | 2390 | 46832 | 32446 | 9337 | **PC O-42:5** | 9 | 6 | 527 | 1101 |
| DG 30:0 | 876062 | 690091 | 3412 | 1211 | **PE O-34:2** | 19 | 40 | 2379 | 2204 |
| DG 32:0 | 15108 | 10355 | 16283 | 7909 | **PE O-38:7** | 3965 | 818 | 8255 | 3819 |
| DG 33:3 | 494939 | 425118 | 489887 | 353403 | **PE O-38:6** | 6714 | 1867 | 7506 | 3016 |
| DG 34:0 | 59157 | 51973 | 51453 | 17344 | **PG O-31:2** | 56486 | 18313 | 9078 | 321 |
| DG 34:1 | 245626 | 151524 | 148212 | 79664 | **PG O-32:2** | 22727 | 3506 | 1511 | 28 |
| DG 34:0 | 102227 | 88615 | 97105 | 37399 | **PG O-34:3** | 597686 | 243078 | 93233 | 2263 |
| DG 35:2 | 26478 | 9386 | 983 | 69 | **PG O-35:3** | 1864 | 135 | 30 | 0 |

| | | | | | | | | | |
|---|---|---|---|---|---|---|---|---|---|
| DG 36:2 | 36421 | 5544 | 11101 | 1467 | **PG O-36:3** | 7562 | 923 | 231 | 30 |
| DG 36:2 | 12809 | 450 | 5591 | 8798 | **PG O-37:3** | 10901 | 1605 | 242 | 46 |
| DG 38:4 | 469768 | 342568 | 371884 | 237228 | **PG O-37:1** | 1150 | 695 | 1783 | 498 |
| DG 39:8 | 211845 | 129385 | 86425 | 42260 | **FA 15:4** | 754 | 422 | 978 | 664 |
| DG 39:7 | 750 | 253 | 22689 | 529 | **FA 16:0** | 6762 | 3743 | 8406 | 4551 |
| DG 41:11 | 99450 | 114098 | 77080 | 30437 | **FA 16:0** | 1141603 | 1228857 | 1661017 | 1338785 |
| DG 43:5 | 355832 | 253494 | 273415 | 156978 | **FA 16:0** | 6567 | 5976 | 7918 | 3011 |
| DG 43:5 | 56658 | 45978 | 53681 | 30205 | **FA 16:0** | 1675470 | 1438111 | 1645772 | 951107 |
| DG 47:13 | 6271185 | 870062 | 220438 | 1469 | **FA 17:1** | 9504 | 7780 | 4224 | 9943 |
| DG 48:13 | 638115 | 62844 | 6279 | 400 | **FA 17:0** | 7233 | 7950 | 12523 | 4191 |
| DG 48:12 | 4352 | 6510 | 14674 | 13322 | **FA 18:2** | 3348 | 9106 | 5505 | 6134 |
| DG 48:11 | 468137 | 150523 | 79551 | 670 | **FA 18:1** | 7971 | 4782 | 9323 | 5538 |
| DG 50:14 | 15527 | 30021 | 15528 | 18 | **FA 18:1** | 7198 | 15037 | 57701 | 6616 |
| DGGA 33:8 | 153648 | 127919 | 727 | 115098 | **FA 18:1** | 221115 | 844629 | 1071083 | 161094 |
| LPE O-22:1 | 17215 | 16251 | 16597 | 16686 | **FA 18:1** | 9493 | 7510 | 10269 | 3759 |
| PC O-29:3 | 204314 | 282455 | 190214 | 124542 | **FA 18:0** | 3256 | 2469 | 4731 | 3176 |
| PC O-30:7 | 9925 | 2807 | 180891 | 73132 | **FA 18:0** | 3184 | 3204 | 5021 | 2273 |
| PC O-30:0 | 89 | 1146 | 78692 | 30866 | **FA 18:0** | 589689 | 715098 | 947363 | 559728 |
| PC O-35:6 | 319 | 161 | 39395 | 22011 | **FA 19:1** | 1497 | 1094 | 2407 | 1519 |
| PC O-37:7 | 20 | 364 | 259050 | 86068 | **FA 19:1** | 5479 | 6033 | 139 | 5531 |
| PC O-39:10 | 374 | 90 | 21316 | 100 | **FA 19:1** | 4218 | 3567 | 647 | 4133 |
| PC O-38:1 | 8998 | 4783 | 4134 | 3039 | **FA 19:0** | 3779 | 15590 | 11414 | 22369 |
| PC O-40:8 | 1062 | 506 | 281236 | 57263 | **FA 19:0** | 6381 | 22841 | 14048 | 24244 |
| PC O-41:10 | 130 | 58 | 27471 | 135 | **FA 20:4** | 37 | 101 | 4577 | 18888 |

| | | | | | | | | | |
|---|---|---|---|---|---|---|---|---|---|
| PC O-41:10 | 58 | 26 | 226 | 2388 | **FA 20:3** | 4417 | 1470 | 4985 | 18366 |
| PC O-40:2 | 61800 | 30053 | 58082 | 23102 | **FA 20:2** | 861 | 975 | 1240 | 476 |
| PC O-42:6 | 16 | 18 | 57318 | 5045 | **FA 20:1** | 4512 | 5395 | 17748 | 4276 |
| PC O-43:11 | 0 | 38 | 407 | 30 | **FA 20:0** | 3057 | 5055 | 5973 | 2147 |
| PE O-36:2 | 74367 | 57689 | 27019 | 370 | **FA 20:0** | 241 | 978 | 573 | 62 |
| PE P-36:1 | 73 | 55687 | 0 | 12907 | **FA 22:0** | 887 | 1534 | 1199 | 1264 |
| PE O-37:8 | 262135 | 27837 | 4633 | 258 | **FA 24:0** | 2400 | 3098 | 2718 | 1572 |
| TG O-31:0 | 23253 | 15730 | 21041 | 10355 | **FA 24:0** | 1046 | 1420 | 1374 | 666 |
| TG O-32:0 | 20720 | 8643 | 10046 | 7424 | **FA 25:0** | 736 | 962 | 1044 | 346 |
| TG O-35:2 | 34714 | 12728 | 5631 | 705 | **FA 25:0** | 901 | 1115 | 1415 | 771 |
| Hex2Cer 34:1;2O | 27 | 75 | 17947 | 9068 | **FA 26:0** | 1763 | 3629 | 4227 | 882 |
| HexCer 36:8;3O | 129253 | 123848 | 71797 | 939 | **FA 27:0** | 410 | 1014 | 1073 | 146 |
| LDGTS 14:1 | 2573 | 665467 | 2987 | 922 | **FA 28:0** | 1194 | 2871 | 4379 | 801 |
| LDGTS 14:1 | 7776 | 3524293 | 11525 | 3017 | **FA 28:0** | 1750 | 1546 | 1951 | 1420 |
| LDGTS 15:1 | 78033 | 51818 | 66624 | 31946 | **FA 32:0** | 47 | 380 | 1748 | 262 |
| LDGTS 18:2 | 44821 | 47204 | 40966 | 40199 | **FA 34:9** | 6594 | 7043 | 11480 | 6234 |
| LPC 16:0 | 316 | 348 | 483028 | 467872 | **FA 34:9** | 18430 | 5936 | 7671 | 33122 |
| LPC 16:0 | 6615 | 1308 | 543814 | 702430 | **FA 34:9** | 48958 | 4190 | 16011 | 13812 |
| LPC 16:0 | 4726 | 850 | 281495 | 205870 | **FA 34:9** | 21565 | 19564 | 16572 | 17839 |
| LPC 18:1 | 546 | 100 | 524754 | 865911 | **FA 34:0** | 1197 | 1109 | 1691 | 532 |
| LPC 18:0 | 650 | 178 | 69188 | 41697 | **FA 36:9** | 2735 | 4827 | 7747 | 4711 |
| LPC 18:0 | 7113 | 6739 | 2285290 | 1117804 | **FA 36:0** | 1787 | 1178 | 1955 | 337 |

| | | | | | | | | | |
|---|---|---|---|---|---|---|---|---|---|
| LPC 17:2 | 1208699 | 1432329 | 1053733 | 798481 | **FA 38:10** | 501 | 447 | 338 | 132 |
| LPC 17:2 | 1049575 | 1029311 | 938807 | 716951 | **FA 38:10** | 1557 | 690 | 1353 | 539 |
| LPC 20:0 | 1127 | 1307 | 361791 | 332088 | **FA 38:10** | 864 | 2521 | 2372 | 549 |
| LPC 22:0 | 2168 | 614 | 110792 | 67714 | **FA 38:9** | 884 | 51 | 181 | 75 |
| LPC 28:7 | 1568 | 297 | 71670 | 354 | **FA 38:9** | 1097 | 64 | 421 | 132 |
| LPC 28:7 | 566 | 299 | 37147 | 424 | **FA 38:9** | 5560 | 20415 | 18239 | 14271 |
| LPC 28:3 | 136392 | 76867 | 152315 | 91611 | **FA 37:0** | 1947 | 1801 | 2833 | 1005 |
| LPE 28:7 | 240761 | 35274 | 121 | 148 | **FA 37:0** | 1370 | 1421 | 1939 | 444 |
| NAE 17:3 | 89058 | 107032 | 90458 | 84391 | **FA 40:0** | 576 | 600 | 1165 | 53 |
| NAE 17:3 | 73850 | 107766 | 52348 | 80843 | **FAHFA 25:0;O** | 991 | 1489 | 377 | 690 |
| NAE 18:3 | 894 | 2464 | 21201 | 15105 | **FAHFA 26:0;O** | 1673 | 2176 | 1774 | 1282 |
| NAE 20:1 | 15407 | 30848 | 13873 | 12262 | **FAHFA 32:0;O** | 1102 | 931 | 1131 | 352 |
| NAE 21:4 | 8304 | 27487 | 7078 | 401 | **FAHFA 35:5;O** | 13817 | 16176 | 6207 | 10300 |
| NAGly 35:6;2O | 59172 | 42 | 448 | 2237 | **FAHFA 35:4;O** | 1920 | 1878 | 1390 | 338 |
| TG 32:0;2O | 308444 | 330707 | 295250 | 140670 | **FAHFA 36:0;O** | 2006 | 1657 | 3013 | 1351 |
| TG 41:1;1O | 22757 | 19987 | 22287 | 6661 | **FAHFA 38:3;O** | 450 | 243 | 1474 | 647 |
| TG 41:1;2O | 1012 | 583 | 78815 | 17231 | **FAHFA 39:5;O** | 5451 | 5088 | 2198 | 3452 |
| TG 43:1;1O | 8369 | 6963 | 9065 | 5197 | **HBMP 48:2** | 15210 | 568 | 259 | 12 |
| TG 44:1;2O | 514 | 492 | 809 | 444 | **HBMP 48:1** | 524 | 76 | 48 | 6 |
| TG 47:4;1O | 8618 | 5317 | 5697 | 3878 | **HBMP 48:1** | 537 | 147 | 61 | 11 |
| TG 57:8;1O | 250601 | 261774 | 58811 | 4236 | **HBMP 50:12** | 944 | 1821 | 360 | 2437 |
| PC 26:1 | 398 | 256593 | 35529 | 0 | **HBMP 50:2** | 2040 | 200 | 192 | 19 |
| PC 30:0 | 7304 | 1197 | 640195 | 397684 | **HBMP 50:2** | 13831 | 557 | 471 | 11 |

| | | | | | | | | | |
|---|---|---|---|---|---|---|---|---|---|
| PC 31:2 | 80454 | 69147 | 61099 | 39909 | **HexCer 45:9;4O** | 11 | 63 | 10351 | 1908 |
| PC 32:2 | 6652 | 2539 | 108104 | 9976 | **HexCer 42:1;3O** | 60 | 16 | 1008 | 1544 |
| PC 32:1 | 31684 | 7185 | 4022826 | 662278 | **LPE-N (FA)32:0** | 214848 | 56762 | 17016 | 275 |
| PC 32:1 | 22263 | 4034 | 4243993 | 542086 | **LPE-N (FA)36:1** | 185390 | 26030 | 14731 | 1704 |
| PC 32:0 | 1142 | 538 | 3608489 | 1562 | **LPC 16:0** | 415 | 123 | 10075 | 106621 |
| PC 33:1 | 7791 | 2645 | 1606567 | 440441 | **LPC 16:0** | 408 | 25 | 12706 | 84443 |
| PC 33:1 | 0 | 237 | 15 | 93285 | **LPC 18:1** | 21 | 32 | 9056 | 87902 |
| PC 34:2 | 78832 | 12347 | 3811171 | 471360 | **LPC 18:1** | 73 | 32 | 61377 | 27957 |
| PC 34:2 | 74537 | 17286 | 2663642 | 189134 | **LPC 18:1** | 32 | 10 | 73563 | 12951 |
| PC 34:2 | 83644 | 3306 | 768585 | 76325 | **LPC 18:1** | 10 | 5 | 9261 | 88699 |
| PC 34:1 | 262045 | 77763 | 11601 | 7362499 | **LPC 18:0** | 404 | 621 | 15212 | 237882 |
| PC 34:1 | 308 | 141 | 27534200 | 3578 | **LPC 18:0** | 910 | 967 | 85506 | 172552 |
| PC 34:0 | 7271 | 1133 | 719274 | 486791 | **LPC 18:0** | 80 | 95 | 24771 | 57791 |
| PC 34:0 | 11243 | 2477 | 451938 | 345123 | **LPC 18:0** | 79 | 17 | 24205 | 7198 |
| PC 35:1 | 2470 | 463 | 1317934 | 517201 | **LPC 18:0** | 16 | 72 | 17749 | 12596 |
| PC 36:5 | 2590 | 299 | 452842 | 63126 | **LPC 19:0** | 15 | 11 | 4246 | 6655 |
| PC 36:4 | 19013 | 4080 | 1099 | 108 | **LPC 19:0** | 16 | 11 | 8990 | 3853 |
| PC 36:3 | 4969 | 1318 | 946823 | 140457 | **LPC 20:3** | 4 | 52 | 14345 | 3406 |
| PC 36:3 | 14108 | 4116 | 2866754 | 749248 | **LPC 20:1** | 16 | 3 | 5244 | 14668 |
| PC 36:2 | 66413 | 586 | 6111327 | 3451 | **LPC 20:0** | 48 | 5 | 1593 | 5663 |
| PC 36:2 | 395 | 138 | 4204 | 1807 | **MGDG 33:1** | 486 | 2650 | 1582 | 61 |
| PC 36:2 | 115 | 50 | 1508 | 796934 | **MGDG 36:1** | 459 | 201 | 1066 | 180 |

| | | | | | | | | | |
|---|---|---|---|---|---|---|---|---|---|
| PC 36:1 | 19674 | 4420 | 7896375 | 3727004 | **NAE 17:0** | 2475 | 421 | 295 | 116 |
| PC 36:1 | 53011 | 14990 | 4105173 | 2126947 | **Dodecylbenzenesulfonic acid** | 103478 | 21956 | 44924 | 37432 |
| PC 37:1 | 508 | 338 | 271683 | 1815 | **Dodecylbenzenesulfonic acid** | 12871 | 10348 | 12932 | 9923 |
| PC 38:6 | 9467 | 2469 | 1262426 | 279317 | **Dodecylbenzenesulfonic acid** | 5591 | 3929 | 9110 | 4429 |
| PC 38:6 | 430 | 325 | 211952 | 22163 | **Dodecylbenzenesulfonic acid** | 152579 | 40006 | 86259 | 62342 |
| PC 38:6 | 682 | 186 | 198224 | 29487 | **Dodecylbenzenesulfonic acid** | 12526 | 9128 | 12403 | 13617 |
| PC 38:6 | 28690 | 4013 | 6753958 | 1507141 | **Dodecylbenzenesulfonic acid** | 69146 | 56707 | 82062 | 52616 |
| PC 38:5 | 247 | 119 | 1314 | 1435 | **Norethisterone acetate** | 62281 | 21991 | 41112 | 36088 |
| PC 38:4 | 51444 | 90 | 4211 | 1941996 | **Norethisterone acetate** | 120260 | 374783 | 464546 | 420989 |
| PC 38:4 | 1637 | 894 | 1404941 | 290938 | **Norethisterone acetate** | 651199 | 313844 | 392439 | 548416 |
| PC 38:3 | 16933 | 3300 | 2275026 | 862496 | **Norethisterone acetate** | 361426 | 145167 | 280580 | 177156 |
| PC 38:2 | 2416 | 550 | 361415 | 151459 | **Norethisterone acetate** | 13490 | 43489 | 48394 | 11452 |
| PC 38:2 | 2884 | 752 | 323997 | 157151 | **Norethisterone acetate** | 475450 | 274248 | 376932 | 380190 |

| | | | | | | | | | |
|---|---|---|---|---|---|---|---|---|---|
| PC 38:2 | 2642 | 108830 | 27727 | 216 | **Norethisterone acetate** | 63577 | 35201 | 72888 | 46476 |
| PC 39:6 | 1095 | 1296 | 121472 | 15132 | **Norethisterone acetate** | 11955 | 7353 | 24330 | 6555 |
| PC 40:8 | 144 | 222 | 49468 | 1727 | **Norethisterone acetate** | 599768 | 446819 | 771619 | 521063 |
| PC 40:7 | 1972 | 839 | 1205834 | 417376 | **Norethisterone acetate** | 109935 | 25616 | 161324 | 35655 |
| PC 40:6 | 6486 | 4170 | 8851741 | 2137158 | **Norethisterone acetate** | 74976 | 31578 | 64856 | 45022 |
| PC 40:6 | 5101 | 1170 | 775247 | 80268 | **Norethisterone acetate** | 24296 | 15702 | 33881 | 20908 |
| PC 40:5 | 15351 | 2171 | 3684706 | 1084638 | **Norethisterone acetate** | 5992 | 3795 | 7791 | 5556 |
| PC 40:5 | 24921 | 12 | 436 | 187 | **Norethisterone acetate** | 24633 | 15017 | 33447 | 22539 |
| PC 41:7 | 691 | 153 | 54832 | 13143 | **Norethisterone acetate** | 270511 | 94002 | 321636 | 116378 |
| PC 41:5 | 482 | 93 | 61198 | 15740 | **Norethisterone acetate** | 126099 | 58302 | 109997 | 84491 |
| PC 42:10 | 46 | 9 | 28050 | 2669 | **Norethisterone acetate** | 24206 | 167743 | 308786 | 291032 |
| PC 40:6 | 817 | 245 | 156 | 93 | **Norethisterone acetate** | 58742 | 298364 | 540209 | 522205 |
| PE 31:8 | 2233 | 39585 | 15416 | 33251 | **Norethisterone acetate** | 102844 | 298924 | 184639 | 246738 |
| PE 30:1 | 413 | 494589 | 208296 | 31 | **Norethisterone acetate** | 237224 | 145228 | 233275 | 316763 |
| PE 30:1 | 1623314 | 297988 | 88886 | 526 | **Norethisterone acetate** | 584368 | 269002 | 243631 | 503202 |

| | | | | | | | | | |
|---|---|---|---|---|---|---|---|---|---|
| PE 30:0 | 5507281 | 901033 | 253147 | 13603 | **Norethisterone acetate** | 76202 | 62498 | 98757 | 94401 |
| PE 31:6 | 102544 | 12071 | 5011 | 976 | **Norethisterone acetate** | 14233 | 48466 | 94455 | 22399 |
| PE 31:1 | 1740993 | 808 | 176577 | 68 | **Norethisterone acetate** | 28806 | 153919 | 283373 | 92898 |
| PE 32:2 | 2353160 | 463962 | 213824 | 660 | **Norethisterone acetate** | 13707 | 69077 | 146735 | 50349 |
| PE 32:1 | 23991276 | 7577914 | 5385720 | 41174 | **Norethisterone acetate** | 29625 | 211862 | 383031 | 177344 |
| PE 33:2 | 1897 | 17027 | 288 | 99 | **Norethisterone acetate** | 200130 | 323995 | 512677 | 704357 |
| PE 33:2 | 3258570 | 785225 | 444657 | 2932 | **Norethisterone acetate** | 378127 | 158094 | 131862 | 176874 |
| PE 33:1 | 19230960 | 8539750 | 4040744 | 53352 | **Norethisterone acetate** | 533095 | 241853 | 217172 | 160589 |
| PE 34:2 | 2886427 | 673426 | 290948 | 1695 | **Norethisterone acetate** | 629648 | 304249 | 560804 | 521321 |
| PE 34:2 | 10705014 | 5575312 | 2647590 | 19733 | **Norethisterone acetate** | 613082 | 286607 | 483999 | 464105 |
| PE 34:2 | 226740 | 42593 | 509 | 247 | **Norethisterone acetate** | 545440 | 329521 | 544718 | 491760 |
| PE 35:2 | 14826520 | 4088456 | 1860994 | 18542 | **FA 16:1;(2OH)** | 485311 | 410820 | 269935 | 370718 |
| PE 35:1 | 6239184 | 846186 | 307009 | 6359 | **FA 16:1;(2OH)** | 242246 | 204102 | 226450 | 101760 |
| PE 36:2 | 18955242 | 4794032 | 2010181 | 19015 | **FA 16:0;(2OH)** | 172603 | 156998 | 105076 | 80725 |
| PE 36:1 | 701165 | 93033 | 39444 | 8858 | **FA 16:0;(2OH)** | 62148 | 62834 | 34413 | 18825 |

| | | | | | | | | | |
|---|---|---|---|---|---|---|---|---|---|
| PG 36:2 | 9723385 | 178 | 288 | 55 | **FA 16:0;(2OH)** | 33590 | 31746 | 46919 | 18582 |
| PG 37:2 | 676643 | 66574 | 34894 | 5096 | **FA 16:2;2O** | 10459 | 10202 | 4158 | 8904 |
| PI 38:4 | 0 | 467 | 0 | 0 | **FA 16:2;2O** | 3741 | 2977 | 3646 | 1199 |
| PI 38:4 | 81 | 1190 | 46132 | 14051 | **FA 16:1;2O** | 97779 | 47312 | 27086 | 35162 |
| SM 32:2;2O | 115 | 14 | 348896 | 18989 | **FA 16:0;2O** | 23829 | 30180 | 11597 | 24936 |
| SM 32:1;2O | 1650 | 629 | 1001974 | 128185 | **FA 18:2;O** | 18418 | 11056 | 11029 | 9590 |
| SM 33:1;2O | 515 | 30863 | 705504 | 162308 | **FA 18:1;O** | 9997 | 8149 | 8279 | 9450 |
| SM 34:2;2O | 1390 | 1110 | 2810062 | 373380 | **FA 18:1;O** | 5618 | 6446 | 3997 | 6312 |
| SM 34:1;2O | 51905 | 21378 | 9534333 | 4089546 | **FA 18:1;O** | 3965 | 3842 | 2197 | 3883 |
| SM 34:1;2O | 19562 | 4940 | 866110 | 354533 | **FA 18:1;(2OH)** | 140994 | 123773 | 36892 | 113348 |
| SM 34:0;2O | 12871 | 2321 | 885048 | 251313 | **FA 18:1;O** | 19031 | 9562 | 9585 | 10552 |
| SM 35:2;2O | 439 | 351 | 87130 | 17444 | **FA 18:1;O** | 5328 | 4738 | 2555 | 4973 |
| SM 36:2;2O | 16 | 371 | 965211 | 484844 | **FA 18:1;O** | 63469 | 60387 | 32242 | 20420 |
| SM 36:1;2O | 15600 | 2124 | 1479017 | 739042 | **FA 18:1;O** | 4040 | 3457 | 1980 | 2958 |
| SM 37:7;3O | 5886087 | 1449675 | 332470 | 2050 | **FA 17:1;2O** | 11777 | 7977 | 4565 | 10804 |
| SM 40:2;2O | 1858 | 2408 | 879 | 1041 | **FA 18:0;O** | 53816 | 48507 | 14117 | 40957 |
| SM 39:2;3O | 4179 | 1781 | 901605 | 624565 | **FA 18:0;(2OH)** | 10233 | 11865 | 6749 | 9306 |
| SM 41:4;2O | 13336 | 9736 | 12969 | 4388 | **FA 18:0;(2OH)** | 36557 | 31508 | 12884 | 28919 |
| SM 40:3;3O | 30 | 586 | 229290 | 161999 | **FA 18:0;(2OH)** | 31298 | 30449 | 11370 | 15338 |
| SM 41:1;2O | 94 | 37 | 2189 | 38416 | **FA 18:2;2O** | 3046 | 2551 | 1231 | 3238 |

| | | | | | | | | | |
|---|---|---|---|---|---|---|---|---|---|
| SM 41:1;2O | 115 | 181 | 171888 | 632 | **FA 18:1;2O** | 11796 | 13329 | 6847 | 4818 |
| SM 41:4;3O | 0 | 280 | 544693 | 273914 | **FA 18:1;2O** | 18375 | 10102 | 10240 | 9279 |
| SM 41:3;3O | 1712 | 528 | 247166 | 183546 | **FA 18:1;2O** | 26170 | 25047 | 8432 | 18930 |
| SM 42:2;2O | 11762 | 365 | 197 | 2092 | **FA 19:0;O** | 2687 | 2242 | 2093 | 387 |
| SM 42:1;2O | 3505 | 23 | 608 | 352575 | **FA 18:0;2O** | 17961 | 15811 | 4618 | 13264 |
| SM 44:6;2O | 160 | 206 | 211 | 8 | **FA 19:1;2O** | 6002 | 8382 | 7101 | 7713 |
| ST 27:1;O | 1225 | 602 | 127978 | 5023 | **FA 20:1;2O** | 1966 | 2227 | 654 | 2303 |
| ST 27:1;O | 1932 | 421 | 82565 | 1295 | **FA 22:0;(2 OH)** | 2062 | 2291 | 2431 | 882 |
| ST 27:1;O | 1989 | 480 | 71283 | 441106 | **FA 22:1;2O** | 4010 | 6050 | 2570 | 3806 |
| TG 26:0 | 340515 | 213421 | 235185 | 62814 | **FA 22:1;2O** | 519 | 501 | 2133 | 116 |
| TG 28:0 | 204693 | 242435 | 225761 | 92101 | **FA 24:0;(2 OH)** | 6515 | 8895 | 8577 | 4550 |
| TG 28:0 | 5043 | 55740 | 54663 | 55440 | **PC 32:0;O** | 26 | 57 | 2746 | 2604 |
| TG 42:1 | 7767 | 8465 | 10373 | 15641 | **PC 33:2;O** | 212 | 85 | 80270 | 15557 |
| TG 42:0 | 16925 | 13855 | 21613 | 24921 | **PC 33:2;O** | 113 | 49 | 174739 | 15727 |
| TG 43:1 | 7493 | 5386 | 6657 | 15042 | **PC 34:1;O** | 38 | 9 | 5201 | 797 |
| TG 43:0 | 10531 | 7299 | 13119 | 19324 | **PC 34:1;O** | 42 | 24 | 24638 | 15254 |
| TG 44:2 | 5411 | 6740 | 8001 | 14962 | **PC 32:1;3O** | 15 | 12 | 18150 | 1035 |
| TG 44:1 | 16606 | 12210 | 19101 | 34826 | **PC 33:1;3O** | 27 | 64 | 4711 | 1101 |
| TG 44:0 | 22365 | 14494 | 24212 | 37022 | **PC 33:1;3O** | 9 | 56 | 9541 | 2526 |
| TG 45:2 | 544 | 5371 | 60 | 17003 | **PC 34:0;2O** | 13 | 26 | 3247 | 173 |
| TG 45:1 | 14564 | 12939 | 19332 | 38975 | **PC 34:1;3O** | 47 | 172 | 668053 | 10255 |
| TG 45:0 | 20010 | 14483 | 23822 | 41587 | **PC 34:0;3O** | 32 | 105 | 14178 | 8806 |

| | | | | |
|---|---|---|---|---|
| TG 46:3 | 112 | 92 | 52 | 9948 |
| TG 46:2 | 17317 | 12162 | 18231 | 38369 |
| TG 47:2 | 16879 | 11118 | 15894 | 35505 |
| TG 47:1 | 27629 | 17786 | 28519 | 67522 |
| TG 48:1 | 36550 | 19594 | 38860 | 101405 |
| TG 44:1;O2 | 162177 | 108020 | 150298 | 66684 |
| VAE 24:3 | 6647 | 22302 | 23017 | 13094 |

| | | | | |
|---|---|---|---|---|
| **PC 38:7;O** | 9 | 27 | 1584 | 190 |
| **PC 38:7;O** | 200 | 46 | 12125 | 719 |
| **PC 38:7;O** | 17 | 18 | 9675 | 10645 |
| **PC 35:1;3O** | 28 | 23 | 30851 | 1642 |
| **PC 38:5;O** | 4 | 17 | 3656 | 1190 |
| **PC 36:4;3O** | 0 | 21 | 12663 | 499 |
| **PC 36:3;3O** | 31 | 10 | 13367 | 2833 |
| **PC 39:8;O** | 15 | 42 | 14142 | 6732 |
| **PC 36:3;3O** | 46 | 28 | 4790 | 7841 |
| **PC 36:3;3O** | 38 | 69 | 2888 | 2092 |
| **PC 36:2;3O** | 43 | 20 | 37727 | 6424 |
| **PC 39:6;O** | 62 | 90 | 17057 | 7646 |
| **PC 39:5;O** | 66 | 4 | 22456 | 9767 |
| **PC 36:0;3O** | 20 | 29 | 17993 | 11929 |
| **PE 25:1;O** | 55307 | 6955 | 1460 | 148 |
| **PE 26:0;O** | 3930 | 20 | 97 | 15 |
| **PE 27:2;O** | 24111 | 3019 | 839 | 11 |
| **PE 27:1;O** | 65099 | 8625 | 2514 | 69 |
| **PE 26:1;2O** | 2677 | 198 | 58 | 0 |
| **PE 28:2;O** | 26830 | 4055 | 1077 | 58 |
| **PE 27:1;2O** | 5627 | 705 | 152 | 20 |
| **PE 27:1;2O** | 6743 | 1498 | 250 | 26 |
| **PE 28:0;O** | 3470 | 371 | 96 | 14 |

| | | | | |
|---|---|---|---|---|
| **PE 27:0;2O** | 4240 | 549 | 177 | 42 |
| **PE 29:2;O** | 57059 | 7348 | 2326 | 39 |
| **PE 29:2;O** | 37677 | 4489 | 1988 | 24 |
| **PE 29:1;2O** | 3366 | 527 | 205 | 35 |
| **PE 30:0;O** | 7490 | 1087 | 388 | 36 |
| **PE 30:0;O** | 9816 | 1133 | 360 | 6 |
| **PE 31:1;O** | 32776 | 4607 | 1296 | 7 |
| **PE 31:1;O** | 112 | 120 | 86 | 0 |
| **PE 31:1;O** | 200 | 12 | 15 | 0 |
| **PE 31:1;O** | 327 | 197 | 59 | 10 |
| **PE 31:1;O** | 580 | 957 | 827 | 78 |
| **PE 32:2;O** | 5948 | 952 | 86 | 8 |
| **PE 32:1;O** | 139747 | 19959 | 7056 | 70 |
| **PE 32:1;O** | 10704 | 574 | 70 | 17 |
| **PE 32:1;O** | 68491 | 9432 | 3054 | 26 |
| **PE 32:0;O** | 2034 | 201 | 99 | 0 |
| **PE 33:2;O** | 7517 | 10116 | 6165 | 59 |
| **PE 33:2;O** | 8936 | 1664 | 518 | 0 |
| **PE 33:1;O** | 30731 | 10025 | 4806 | 13 |
| **PE 34:2;O** | 60431 | 8983 | 3094 | 21 |
| **PE 34:2;O** | 75663 | 12650 | 5053 | 48 |
| **PE 34:1;O** | 35392 | 11078 | 1725 | 27 |
| **PE 34:1;O** | 81895 | 12445 | 4812 | 53 |

| | | | | |
|---|---|---|---|---|
| **PE 32:0;3O** | 18 | 42 | 2889 | 1417 |
| **PE 35:3;O** | 4894 | 7375 | 3968 | 12 |
| **PE 35:3;O** | 9688 | 4385 | 1057 | 10 |
| **PE 35:2;O** | 710 | 8587 | 4380 | 45 |
| **PE 34:2;2O** | 15760 | 3831 | 108 | 59 |
| **PE 34:1;2O** | 3415 | 202 | 45 | 8 |
| **PE 36:2;O** | 58249 | 8583 | 3232 | 33 |
| **PE 36:2;O** | 6206 | 553 | 123 | 9 |
| **PE 34:0;3O** | 38 | 54 | 5272 | 1402 |
| **PE 34:0;3O** | 24 | 59 | 5911 | 5282 |
| **PC 30:0** | 29 | 103 | 41117 | 22747 |
| **PC 31:0** | 0 | 31 | 22066 | 14167 |
| **PC 31:0** | 54 | 103 | 934 | 1271 |
| **PC 32:2** | 214 | 84 | 2631 | 71 |
| **PC 32:2** | 167 | 176 | 12204 | 1581 |
| **PC 32:1** | 39 | 70 | 47420 | 2730 |
| **PC 32:1** | 62 | 89 | 142780 | 57260 |
| **PC 32:1** | 322 | 49 | 60538 | 31279 |
| **PC 32:0** | 31 | 83 | 7818 | 5394 |
| **PC 32:0** | 45 | 87 | 70383 | 22692 |
| **PC 32:0** | 16 | 176 | 56862 | 66078 |
| **PC 32:0** | 45 | 0 | 2668 | 5803 |
| **PC 32:0** | 61 | 38 | 2175 | 14441 |
| **PC 32:0** | 122 | 71 | 1272 | 5645 |
| **PC 33:1** | 66 | 37 | 58878 | 52730 |
| **PC 33:0** | 135 | 39 | 11852 | 4621 |
| **PC 34:4** | 14 | 142 | 4278 | 240 |
| **PC 34:2** | 123 | 202 | 61871 | 3586 |
| **PC 34:2** | 84 | 340 | 38976 | 5325 |
| **PC 34:2** | 77 | 300 | 273594 | 85356 |
| **PC 34:2** | 96 | 168 | 23862 | 26126 |
| **PC 34:1** | 329 | 345 | 343774 | 191847 |
| **PC 34:1** | 370 | 181 | 428479 | 256283 |
| **PC 34:1** | 124 | 168 | 63262 | 19846 |
| **PC 34:1** | 627 | 1836 | 131908 | 210922 |
| **PC 34:1** | 166 | 165 | 315872 | 51345 |

| | | | | |
|---|---|---|---|---|
| **PC 34:1** | 489 | 319 | 271734 | 703704 |
| **PC 34:1** | 321 | 255 | 39037 | 27677 |
| **PC 34:0** | 193 | 162 | 64722 | 47955 |
| **PC 35:4** | 105 | 20 | 3784 | 178 |
| **PC 35:1** | 61 | 75 | 13069 | 38605 |
| **PC 35:1** | 107 | 80 | 142832 | 77474 |
| **PC 36:5** | 0 | 40 | 3810 | 1290 |
| **PC 36:5** | 0 | 14 | 36778 | 9565 |
| **PC 36:4** | 59 | 71 | 388800 | 8683 |
| **PC 36:4** | 86 | 129 | 394934 | 119502 |
| **PC 36:3** | 62 | 316 | 273148 | 99430 |
| **PC 36:2** | 145 | 210 | 393559 | 12773 |
| **PC 36:2** | 172 | 187 | 87023 | 151173 |
| **PC 36:2** | 699 | 855 | 227489 | 104051 |
| **PC 36:1** | 715 | 199 | 213462 | 449969 |
| **PC 36:1** | 16 | 26 | 1440 | 331 |
| **PC 36:1** | 1492 | 233 | 35320 | 17539 |
| **PC 36:1** | 3271 | 546 | 39381 | 152194 |
| **PC 37:6** | 0 | 20 | 11639 | 2349 |
| **PC 37:5** | 0 | 34 | 3732 | 1615 |
| **PC 37:4** | 21 | 22 | 1388 | 760 |
| **PC 37:4** | 63 | 5 | 5202 | 2352 |
| **PC 37:3** | 19 | 9 | 4539 | 3052 |
| **PC 38:6** | 69 | 286 | 194327 | 166561 |
| **PC 38:5** | 77 | 53 | 10156 | 3782 |
| **PC 38:5** | 36 | 97 | 131261 | 46475 |
| **PC 38:5** | 155 | 5 | 223340 | 198921 |
| **PC 38:4** | 60 | 99 | 169666 | 16717 |
| **PC 38:4** | 136 | 298 | 658197 | 270265 |
| **PC 38:4** | 54 | 29 | 38786 | 33308 |
| **PC 38:4** | 0 | 48 | 15338 | 6802 |
| **PC 38:4** | 31 | 77 | 16845 | 31077 |
| **PC 38:4** | 40 | 71 | 84234 | 2807 |
| **PC 38:3** | 94 | 95 | 110658 | 46849 |
| **PC 38:3** | 19 | 24 | 245416 | 111708 |
| **PC 38:3** | 19 | 23 | 141262 | 66048 |
| **PC 38:3** | 155 | 155 | 30653 | 25203 |
| **PC 38:2** | 84 | 18 | 7965 | 13232 |
| **PC 38:2** | 75 | 68 | 9884 | 15281 |
| **PC 38:1** | 12 | 7 | 2661 | 1646 |
| **PC 38:1** | 10 | 1 | 2794 | 6369 |
| **PC 39:5** | 74 | 3 | 6817 | 8695 |
| **PC 39:4** | 17 | 5 | 2390 | 978 |
| **PC 40:6** | 52 | 32 | 23553 | 24592 |
| **PC 40:6** | 59 | 135 | 250156 | 61243 |
| **PC 40:6** | 53 | 146 | 134543 | 193415 |

| | | | | |
|---|---|---|---|---|
| **PC 40:6** | 16 | 16 | 8328 | 1495 |
| **PC 40:6** | 98 | 18 | 24394 | 21793 |
| **PC 40:5** | 43 | 64 | 411338 | 10541 |
| **PC 40:5** | 202 | 88 | 101393 | 94521 |
| **PC 40:5** | 84 | 283 | 19889 | 71028 |
| **PC 40:5** | 52 | 122 | 29636 | 11060 |
| **PC 40:3** | 11 | 17 | 2726 | 284 |
| **PC 41:9** | 22 | 1 | 1694 | 260 |
| **PC 41:6** | 36 | 18 | 5906 | 1476 |
| **PE 26:1** | 8963 | 890 | 460 | 100 |
| **PE 26:0** | 21053 | 2628 | 583 | 12 |
| **PE 28:1** | 45192 | 7203 | 2303 | 14 |
| **PE 28:0** | 260023 | 32807 | 12277 | 45 |
| **PE 29:1** | 24732 | 4746 | 1565 | 24 |
| **PE 29:0** | 10763 | 3315 | 915 | 22 |
| **PE 30:1** | 877781 | 151931 | 59586 | 42 |
| **PE 30:1** | 423381 | 74938 | 29777 | 255 |
| **PE 30:0** | 760646 | 245866 | 19428 | 792 |
| **PE 31:2** | 12084 | 2088 | 739 | 8 |
| **PE 31:1** | 532319 | 96106 | 42841 | 272 |
| **PE 31:0** | 17082 | 3639 | 1558 | 51 |
| **PE 32:2** | 554525 | 180811 | 79065 | 360 |
| **PE 32:2** | 27715 | 5529 | 1707 | 24 |
| **PE 32:1** | 112848 | 18438 | 8138 | 173 |
| **PE 32:1** | 198248 | 97 | 8 | 6 |
| **PE 32:1** | 13590 | 3020 | 813 | 68 |
| **PE 32:1** | 3071406 | 2097433 | 767509 | 12119 |
| **PE 32:1** | 3484 | 506 | 177 | 30 |
| **PE 33:2** | 34633 | 12090 | 2447 | 62 |
| **PE 33:2** | 16409 | 3350 | 1036 | 91 |
| **PE 33:2** | 755482 | 195710 | 77269 | 844 |
| **PE 33:1** | 39458 | 5703 | 2663 | 48 |
| **PE 33:1** | 80509 | 18374 | 7820 | 129 |
| **PE 33:1** | 6919 | 1109 | 765 | 44 |
| **PE 33:1** | 2388867 | 2050750 | 1074355 | 19133 |
| **PE 33:1** | 308950 | 108580 | 40606 | 1657 |
| **PE 34:3** | 9750 | 2487 | 416 | 9 |
| **PE 34:2** | 629720 | 169800 | 81541 | 1141 |
| **PE 34:2** | 1127370 | 1364254 | 623709 | 6149 |
| **PE 34:1** | 574842 | 111231 | 43704 | 2325 |
| **PE 34:1** | 34071 | 4711 | 2189 | 62 |
| **PE 34:1** | 8000 | 1130 | 358 | 47 |
| **PE 34:1** | 7550 | 1292 | 335 | 13 |

| | | | | |
|---|---|---|---|---|
| **PE 34:1** | 2242047 | 1655946 | 729615 | 14095 |
| **PE 34:1** | 5480 | 256 | 340 | 31 |
| **PE 34:0** | 15663 | 2224 | 1058 | 181 |
| **PE 35:2** | 1357722 | 955868 | 361851 | 5398 |
| **PE 35:2** | 43488 | 11128 | 4633 | 30 |
| **PE 35:2** | 15248 | 9495 | 4337 | 196 |
| **PE 35:1** | 1085506 | 235204 | 108251 | 2100 |
| **PE 36:2** | 1828463 | 1139708 | 460405 | 7877 |
| **PE 36:2** | 394267 | 68372 | 34231 | 1042 |
| **PE 37:2** | 554600 | 138969 | 67612 | 1484 |
| **PE 37:1** | 12869 | 1212 | 398 | 28 |
| **PE 38:2** | 24833 | 3937 | 632 | 51 |
| **PE 38:2** | 17394 | 3709 | 1743 | 28 |
| **PE 42:8** | 16 | 112 | 1820 | 311 |
| **PE 42:7** | 0 | 23 | 10427 | 872 |
| **PE 44:10** | 19 | 25 | 3393 | 397 |
| **PG 26:0** | 7817 | 713 | 111 | 13 |
| **PG 28:1** | 13283 | 1365 | 198 | 25 |
| **PG 28:0** | 124229 | 10994 | 1893 | 9 |
| **PG 30:1** | 546999 | 49922 | 10500 | 32 |
| **PG 30:1** | 251670 | 21372 | 4782 | 109 |
| **PG 30:1** | 9296 | 322 | 145 | 26 |
| **PG 30:0** | 561143 | 65425 | 17533 | 133 |
| **PG 30:0** | 488 | 425 | 100 | 18 |
| **PG 31:1** | 161659 | 12678 | 4455 | 28 |
| **PG 31:1** | 74979 | 4297 | 1239 | 19 |
| **PG 31:1** | 12800 | 5374 | 2112 | 39 |
| **PG 32:2** | 25329 | 1826 | 413 | 0 |
| **PG 32:2** | 481350 | 46396 | 11018 | 9 |
| **PG 32:1** | 6040 | 335 | 40 | 31 |
| **PG 32:1** | 2518980 | 988449 | 290174 | 2270 |
| **PG 32:1** | 5879359 | 357223 | 98769 | 439 |
| **PG 32:0** | 18313 | 2559 | 585 | 99 |
| **PG 33:2** | 414190 | 28251 | 9293 | 42 |
| **PG 33:2** | 12227 | 2557 | 205 | 4 |
| **PG 33:2** | 1188 | 2875 | 2093 | 13 |
| **PG 33:1** | 153762 | 8396 | 2183 | 169 |
| **PG 33:1** | 2787442 | 218455 | 9731 | 335 |

| | | | | |
|---|---|---|---|---|
| **PG 34:3** | 22563 | 1551 | 152 | 43 |
| **PG 34:2** | 21567 | 12842 | 454 | 46 |
| **PG 34:2** | 3748333 | 521303 | 151796 | 921 |
| **PG 34:2** | 357265 | 34498 | 8037 | 136 |
| **PG 34:2** | 78750 | 8303 | 818 | 53 |
| **PG 34:1** | 2927334 | 201553 | 40476 | 741 |
| **PG 34:1** | 9277 | 124 | 80 | 10 |
| **PG 34:1** | 10091 | 2595 | 679 | 44 |
| **PG 34:1** | 21782 | 2448 | 278 | 17 |
| **PG 34:1** | 29082 | 2645 | 392 | 63 |
| **PG 34:1** | 16315 | 484 | 94 | 20 |
| **PG 34:1** | 1494894 | 1279182 | 216228 | 4281 |
| **PG 34:1** | 4117 | 211 | 151 | 4 |
| **PG 35:2** | 1014697 | 131006 | 41136 | 310 |
| **PG 35:1** | 587769 | 27657 | 6472 | 182 |
| **PG 36:3** | 16819 | 1639 | 217 | 7 |
| **PG 36:2** | 3547544 | 803465 | 273107 | 3791 |
| **PG 36:2** | 2361 | 221 | 52 | 8 |
| **PG 36:2** | 1597 | 56 | 9 | 14 |
| **PG 36:2** | 23487 | 1249 | 470 | 0 |
| **PG 36:2** | 8603 | 4460 | 579 | 15 |
| **PG 36:2** | 86916 | 14466 | 1499 | 34 |
| **PG 36:2** | 4884 | 83 | 120 | 9 |
| **PG 36:1** | 219620 | 18350 | 3223 | 138 |
| **PG 37:2** | 89154 | 4016 | 6812 | 43 |
| **PG 37:2** | 125820 | 35691 | 9989 | 134 |
| **PG 37:2** | 9738 | 1619 | 193 | 17 |
| **PG 37:2** | 1698 | 805 | 225 | 85 |
| **PG 37:2** | 1610 | 1054 | 493 | 37 |
| **PG 37:2** | 1752 | 853 | 481 | 38 |
| **PG 37:1** | 12910 | 162 | 270 | 18 |
| **PG 38:2** | 11835 | 1708 | 191 | 23 |
| **PG 38:2** | 5298 | 223 | 200 | 4 |
| **PG 38:2** | 19885 | 2605 | 594 | 22 |
| **PG 42:10** | 7480 | 1013 | 646 | 25 |
| **SM 34:1;2O** | 2578 | 2595 | 1692251 | 911255 |
| **SM 34:0;2O** | 36 | 197 | 40345 | 37607 |

| | | | | |
|---|---|---|---|---|
| **SM 34:2;3O** | 37 | 13 | 39349 | 9363 |
| **SM 35:1;2O** | 114 | 69 | 47765 | 28444 |
| **SM 36:1;2O** | 171 | 135 | 59668 | 169403 |
| **SM 38:1;2O** | 122 | 95 | 33607 | 40315 |
| **SM 41:7;2O** | 272 | 141 | 161635 | 119993 |
| **SM 41:1;2O** | 90 | 17 | 13470 | 39441 |
| **SM 42:3;2O** | 360 | 163 | 41072 | 89416 |

**Extended Data Movie 1: Density map of the P116 homodimer**

Rotation along the long axis of the cryoEM density map. The complete extracellular region of the P116 dimer at 3.3 Å resolution, 90 degrees apart. The individual domains can be appreciated: The dimerization interface (shown in pink), the core domains with the four contiguous antiparallel helices (shown in blue) and a β-sheet with five antiparallel strands (shown in orange) as well as the N-terminal domain is shown in green (Color coding is identical to the coloring in the Figures 1 &2).

**Extended Data Movie 2: Ribbon model of P116**

Ribbon representation of the P116 structure with the same coloring as in Movie 1

**Extended Data Movie 3: Flexibility of P116**

Animation of the wringing motion between the monomers as seen after classification of the cryoEM data. In some classes both cavities face the same direction, while in other they face 80 degrees apart. The speed of the movie may be adjusted in order to appreciate the conformational change. The N- terminal domains were computationally removed for the better visualization.

**Extended Data Movie 4: Hydrophobicity map of the P116 homodimer**

Rotation along the long axis of the hydrophobic map of P116. The huge hydrophobic cavity that is fully accessible to solvent can be appreciated.

**Extended Data Movie 5: Crosssection of one P116 monomer**

Cross-section of the hydrophobic map of the monomer reveals the positions of the ligands (in red).

**Extended Data Movie 6: Ribbon model of a P116 monomer (colors as in Figure 2 & 3) with the unaccounted elongated densities.**

Ribbon representation of the monomer with the ligands (in red). The alignment of the ligands along the bridge helix can be seen.

**Extended Data Movie 7: Conformational change of P116 top view**

Morphing between the ribbon models from P116 and P116 empty. Conformational changes of the P116 shown from the top view. Starting from the structure of the full P116 (open) a morphing is shown towards the structure of the empty P116. The speed of the movie may be adjusted in order to appreciate the conformational change.

**Extended Data Movie 8: Conformational change of P116 distal view**
Morphing between the ribbon models from P116 and P116 empty. Conformational changes of the P116 shown from an arbitrary view. The movement of the four fingers (in blue) towards the core domain (in orange) can be appreciated. The speed of the movie may be adjusted in order to appreciate the conformational change.

**Extended Data Movie 9: Conformational change of P116 distal view with ligands**
Morphing between the ribbon models from P116 and P116 empty. Similar view as in movie 8, only the ligands (in red) present. In the closed conformation the 4 fingers (in blue) clash with the ligands.
